# Potential mode of action of multispecies inoculums on wheat growth under water stress

**DOI:** 10.1101/2025.02.18.638906

**Authors:** Asmaâ Agoussar, Julien Tremblay, Étienne Yergeau

**Affiliations:** Centre Armand-Frappier Santé Biotechnologie, Institut national de la recherche scientifique, 531 boulevard des Prairies, Laval, QC, H7V 1B7, Canada

## Abstract

Manipulating microbial communities could increase crop resistance to environmental stressors such as drought. It is, however, not clear what would be the best approach to do so and what microbial traits are important. Here, we first compare multispecies inoculums created using different approaches. The only inoculum that increased wheat fresh biomass under drought was the one created from 25 isolates that had showed a capacity to grow under high osmolarity. We then looked at two potential mechanisms of action of this inoculum: 1) direct action, by sequencing and screening the genomes of the inoculated bacteria, 2) indirect action, by sequencing the 16S rRNA gene and ITS region of rhizosphere, root and leaves microbial communities. The microbes in the inoculum harbored many traits related to plant growth promoting, competition and water stress resistance. The inoculation also resulted in significant shifts in the microbial communities associated with wheat, including some microorganisms previously reported to improve plant drought resistance. We conclude that the inoculum studied here increased wheat growth because it potentially acted on two fronts: directly, through the traits it was selected for, and indirectly, through inducing shifts in the resident plant microbial communities.

## Introduction

Manipulating or engineering the microbiota could help crops better resist to drought [1–3]. Some microorganisms can increase the resistance of wheat to environmental stress [4–6]. Conversely, changes in soil water availability also affected the microbial communities associated with wheat [7–9,9–13]. In one of our studies, we observed that most changes in the wheat microbiota under water stress resulted from alterations in the relative abundance of already present microbes, with little recruitment from a drought-adapted multispecies inoculum or from the bulk soil [9]. This might be due to modifications in the plant exudation patterns under water stress [14] but might also be an indirect effect of the inoculation. The inoculated strains can modify the resident microbial communities, either by changing the community structure [15,16] or its functional potential by horizontal gene transfer [17–19]. This could indirectly impact plant growth and resistance to water stress, on top of the direct effect of the inoculated microorganisms.

The inoculation of mixed microbial isolates can directly enhance crop growth in stressful conditions [20–22]. To enhance growth under water stress, the inoculated microbes should not only be able to resist to water stress but also to promote plant growth, through provision of nutrients and manipulation of plant hormones, among others [23–25]. Microorganisms can also increase plant drought resistance [26,27], through manipulations of plant hormones [28,29], provision of osmolytes [30–34] or modification of the plant epigenetics [35,36]. Additionally, the production of secondary metabolites could enhance the competitive ability of the microorganisms, increasing the chance they will persist in the plant environment after inoculation. Clearly, a wide repertoire of traits is necessary for a microorganism to establish in the plant environment under drought and change its host phenotype.

Another factor to consider is the compatibility of the inoculated microorganisms with the host – this will define their capacity to colonize and establish themselves in the plant environment. For instance, Arabidopsis plants grown in their native soil exhibited greater resistance to moderate drought than those grown in soil where corn or pine was grown [37]. Microbial communities also vary from one plant compartment to another [8,38], resulting in different capacities to grow at low water availabilities [38]. A better understanding of the abovementioned direct and indirect mechanisms is crucial for the rational design of multi-species inoculum.

Multi-species inoculum can be created using various approaches [39]. The traditional isolation-screening approach is used to select the isolates with the most suitable traits. Although some studies have demonstrated the efficiency of this approach for inoculum creation [20,40,41], comparison of approaches and in-depth characterization beyond the plant responses are seldom performed. Here, we compare the effects of four different multi-species inoculums on wheat growth under normal and water stress conditions. Furthermore, we sequenced the genomes of the isolates used to develop the most effective inoculum and screened them for key traits related to life in the plant environment under water stress. We also examined the impact of the inoculation on the wheat-associated bacterial and fungal communities. This analysis provides insights into the potential mechanisms underlying the effects of multi-species inoculum on wheat growth under water stress.

## Material and methods

### Inoculum preparation

#### Enrichment approach

Two inoculums were prepared based on the community enrichment approach [39]. Soil from our experimental field at the at the Armand-Frappier Santé Biotechnologie Centre (Laval, Québec, Canada) was incubated at 30°C for two months under either 1) dry conditions, in an aluminum plate without cover (“Enrich-Dry” inoculum), or 2) moist conditions using a plastic box with a cover (“Enrich-Moist” inoculum). The moist soil was soaked with sterile water each week. The extraction of the soil microorganisms was done as previously described [42].

#### Isolation-based approach

Two inoculums were prepared with a mixture of 25 isolates each. The isolation, identification and screening of the microorganisms for their capacity to grow under high osmotic pressure (30% PEG) were done in Agoussar et al. [38]. Here, the isolates were evaluated for their potential to promote wheat germination and growth. We sterilized the surface of wheat seeds by soaking them in 70% Ethanol for 2 min and 0.5% NaOCl for 2 min, followed by five washings in sterile water. We measured the percentage of germination after three days of incubation at room temperature in sterile petri dishes following inoculation with the different isolates vs. a sterile media control. We planted surface sterilized seeds in sterile soil in closed Magenta boxes before inoculating them with the microbial isolates or a medium control. The boxes were placed at room temperature and watered daily with sterile water for one week. We categorized the germination and growth promotion capacities as positive, neutral or negative, by comparing them with the uninoculated controls. We ended up selecting 25 isolates (23 bacteria and 2 fungi) that were able to grow under high osmotic pressure and showed positive results for germination and growth promotion assays (“Screening” inoculum). We also created a control inoculum (“Random” inoculum) by randomly selecting 25 isolates from the 542 isolates of Agoussar et al. [38].

Both isolate-based inoculums were prepared by mixing 1 ml (10^4^ CFU/ml) of each isolate growing in a liquid media (TSB for bacteria and YPD for fungal). The resulting 25 ml was centrifuged and cells were resuspended in sterile potassium phosphate buffer and then maintained in 25% glycerol at 4°C.

#### Commercial inoculum

We used, as positive control, a commercial biofertilizer containing a mixture of *Mycorrhiza spp., Trichoderma spp*. and *Bacillus spp*. said to improve wheat growth under drought conditions. As per the provided instructions, we centrifuged 40 μl per 10 g of wheat seeds. As for the other inoculums, the pellet was resuspended with 160 μl of sterile potassium phosphate buffer and maintained in 25% glycerol at 4°C.

#### Negative control

Our negative control contained only a sterile potassium phosphate buffer and 25 % glycerol. All five inoculums and the negative control were stored at 4°C until use.

### Plant growth experiment

The soil used in this experiment was collected from our experimental field, mixed with 1/3 of sand, sieved at 2 mm and autoclaved each 24 h for three successive days. Wheat seeds (*Triticum aestivum)* were surface sterilized as described by Tardif et al. [43] and soaked in the different inoculum for one to two hours until being seeded in the autoclaved soil. We used five seeds per pot, and three days after germination we thinned them to three plantlets per pot.

We grew the plants in a growth cabinet at 70% humidity for a cycle of 6 h of darkness at 21°C, 1h of transition at intermediate level of luminosity at 21°C, 16 h of high luminosity at 25°C, and then 1 h of transition in intermediate level of luminosity at 21°C, for four weeks. The experiment was performed in four different growth chambers, which were considered as experimental blocks for the statistical analyses, with twelve randomly placed pots per growth chamber (two per inoculum). We watered all the pots for the first two weeks. For the next two weeks, six pots per growth chamber (one per inoculum) were maintained in normal conditions at 50% of soil water holding capacity (SWHC), while the remaining pots were maintained at 15% SWHC. We supplemented each pot with 1 ml per week of Hoagland’s nutrient solution [44].

The number of leaves was recorded weekly. At the end of the four-week growth period, samples of roots, leaves, and rhizosphere were collected. For each sample, the length and weight of roots and leaves were measured, and the total fresh and dry weight of the plants was determined. The foliar water content was calculated as follows: (leaves fresh weight - leaves dry weight) / leaves dry weight * 100. The fresh weight was measured immediately after harvest, and the dry weight was obtained after incubating the leaves at 65°C for 72 hours. Based on the plant growth results obtained after four weeks, samples from the best performing inoculum (“Screening”) were selected together with the negative control for further analysis (total of 16 pots: 4 replicates x 2 treatments x 2 water availability).

### DNA extraction and sequencing

Total DNA extraction was performed on 0.5 g of the wheat rhizosphere, roots and leaves samples using the Qiagen kit (DNeasy PowerLyzer PowerSoil) (48 DNA samples: 16 pots x 3 compartments). The library preparation and MiSeq (Illumina) sequencing for 16S rRNA gene and ITS region amplicons using the primers (520F-799R) for 16S and (ITS1F-58A2R) for ITS was performed at the Centre d’expertise et de service Génome Québec (CESGQ, Montréal, Canada). We sequenced the genomes of the 23 bacterial isolates (excluding the two fungal isolates) composing the “Screening” inoculum using PacBio (for most isolates) or MiSeq (Illumina) (two isolates) at the CESGQ.

### Bioinformatic analyses

Amplicon sequencing data (16S rRNA gene and ITS region) were analyzed using AmpliconTagger [45]. Remaining high-quality reads free of sequencing adapters artifacts were dereplicated at 100% identity and clustered/denoised at 99% (DNAclust v3). Clusters of less than three reads were discarded and the remaining clusters were scanned for chimeras using UCHIME, first in de novo mode and then in reference mode [46]. The remaining clusters were clustered at 100% identity (DNAclust v3) to produce ASVs. ASVs were assigned a taxonomic lineage with the RDP classifier using the Silva release 128 databases [47]

The genomic assembly for the 21 bacterial genomes sequenced by the Pacific Biosciences technology was performed at the CESGQ, while the genomic assembly for the two isolates sequenced by MiSeq was performed by SPAdes. The annotation of the bacterial genome was performed based on ‘Prokka’ Prokaryotic genome annotation on Galaxy (Version 1.14.6+galaxy1) (http://bitly.ws/DuAz). Comparative genomic analysis of bacterial genomes was conducted using the OrthoFinder tool on Galaxy (Version 2.5.4+galaxy1) to identify orthogroups within a collection of proteomes and to uncover conserved gene families across the 23 bacterial species. The Diamond research program was employed, and the gene tree inference method was used based on multiple sequence alignment (MSA) using the Muscle program, with a FastTree as a tree inference method. The analysis of secondary metabolite biosynthesis gene clusters in different bacterial isolates was performed using the antiSMASH tool (version: 7.0.0beta2-86685d9d).

### Statistical analysis and data visualization

Statistical analyses were performed in R (version 4.0.3, The R Foundation for Statistical Computing). If the Shapiro-Wilks and Levene tests revealed that, even after log or square root transformation, the alpha diversity and relative abundance data did not meet the assumptions for parametric ANOVA, then independent one-way Kruskal-Wallis tests by rank were performed for the effects of Irrigation, Compartment, Treatment and Block. The effect of Treatment with Screening-SynCom, Compartment and Irrigation on the bacterial and fungal community structure was visualized using principal coordinate analyses (PCoA) and tested using Permanova with 1,000 permutations (including Blocks), both based on Bray-Curtis dissimilarity calculated from the normalized ASV tables. Differential abundance analysis was performed to compare inoculated samples against control samples across three plant compartments under drought and normal conditions. The DESeq2 package in R was used for statistical analysis, applying a negative binomial generalized linear model to account for overdispersion in the count data. Significance thresholds were set at an absolute log2 fold change ≥ 1 and an adjusted p-value (padj) ≤ 0.05, corrected for multiple comparisons using the Benjamini-Hochberg method. The visualization of differentially abundant ASVs was achieved using volcano plots generated with the ggplot2 package in R. The results from plant analyses were averaged over the three plants per pot. When the effect of irrigation was significant, the analysis was then performed for the effect of Compartment, Treatment and Block for the two irrigation regimes separately. The Dunn test was used to compare the effect of the inoculation on germination within the treatments.

## Results

### Plant leaves biomass

Under water stress, the inoculations significantly increased leaves fresh weight (ANOVA, P=0.0261). The plants exposed to the “screening” inoculum had a fresh biomass 67.3% higher than the uninoculated control (Tukey-HSD: P=0.08) (Fig. 1). They were also heavier than the plants exposed to the commercial biofertilizer (Tukey-HSD: P=0.01) and the “enrichment-moist” inoculum (Tukey-HSD: P=0.045) (Fig. 1). As compared to the uninoculated controls, the “screening” inoculum also increased the plant fresh biomass by 24.4% under well-watered conditions and increased dry weight by 46.9% and 111% under well-watered and drought conditions, respectively (Fig. 1). These differences were, however, not statistically significant. At this point, the “screening” inoculum was the best candidate to improve wheat growth under dry and normal conditions and was therefore selected for further analyses.

**Figure 1.**
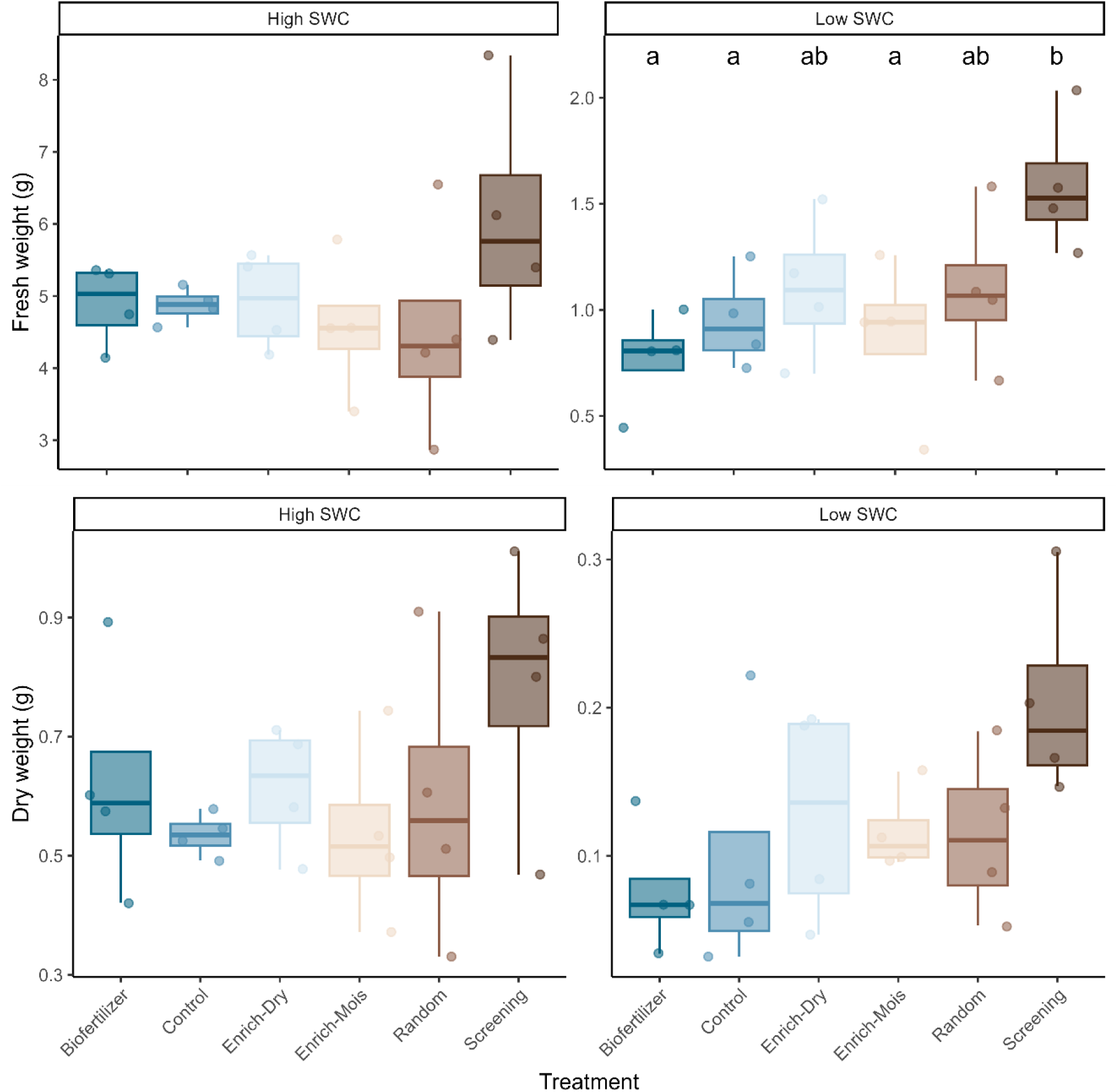
Multispecies inoculums increased plant biomass. Wheat leaves fresh and dry weight following inoculation with five different communities or a buffer control after two weeks of growth under high (50% soil water holding capacity, SWHC) or low (15% SWHC) soil water content (SWC).

### Microbial community structure, composition and diversity following inoculation

We first looked if the “screening” inoculum modified the wheat leaves, roots and rhizosphere microbial communities. As expected, the compartments structured microbial communities, explaining 12.3% (fungi, P=0.001) to 20.8% (bacteria, P=0.001) of the variation observed (Fig. 2 and Table 1). Watering explained a much smaller part of the variation, with 3.8% for bacteria (P=0.002) and a non-significant 2.5% for fungi (P=0.141) (Table 1). The inoculation with the “screening” consortium also significantly influenced the microbial communities, explaining 3.2% of the variation for bacteria (P=0.015) and 4.8% for fungi (P=0.007) (Table 1).

**Figure 2.**
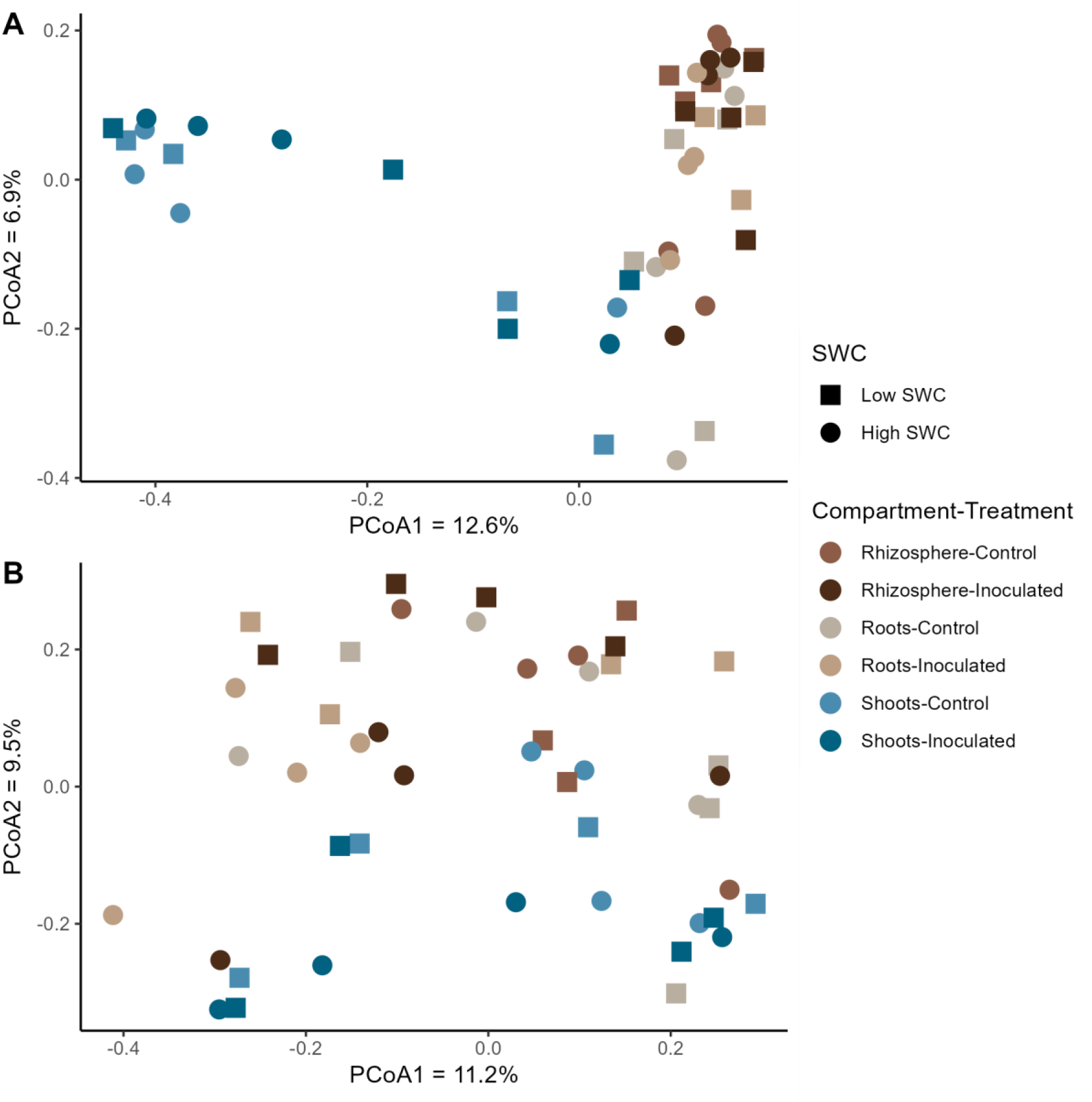
Microbial communities varied according to plant compartment, soil water content and inoculation. Principal coordinates analysis of Bray–Curtis dissimilarities, calculated from the bacterial 16S rRNA gene (a) and the fungal ITS region (b) ASVs tables, for shoot root and rhizosphere samples from wheat inoculated or not with the “screening” inoculum and grown under high (50% soil water holding capacity, SWHC) or low (15% SWHC) soil water content (SWC).

**Table 1.** Permanova tests for the effects of inoculations, plant compartment, soil water content (SWC) and their interactions on the bacterial and fungal community structure based on Bray-Curtis dissimilarities.

|  | Bacteria |  |  | Fungi |  |  |
| --- | --- | --- | --- | --- | --- | --- |
|  | F | R<sup>2</sup> | P | F | R<sup>2</sup> | P |
| Compartment (C) | 6.5949 | 0.208 | **0.001** *** | 3.2493 | 0.123 | **0.001** *** |
| Inoculation (I) | 2.0625 | 0.032 | **0.015** * | 2.5568 | 0.048 | **0.007** ** |
| SWC | 2.4604 | 0.038 | **0.002** ** | 1.3516 | 0.025 | 0.141 |
| block | 2.4732 | 0.117 | **0.001** *** | 1.3912 | 0.079 | **0.054** . |
| C:I | 0.9858 | 0.031 | 0.486 | 0.9189 | 0.034 | 0.573 |
| C:SWC | 0.8477 | 0.026 | 0.724 | 0.8529 | 0.032 | 0.683 |
| I:SWC | 1.3758 | 0.021 | 0.125 | 1.5129 | 0.028 | **0.084** . |
| C:I:SWC | 0.5440 | 0.017 | 0.996 | 0.5294 | 0.020 | 0.987 |

\*\*\*: $P < 0.001$ , \*\*: $0.001 < P < 0.01$ , \* $0.01 < P < 0.05$ , .: $0.05 < P < 0.10$

The inoculation with the “screening” consortium resulted in 9.59%, 26% and 26% increases in bacterial Shannon diversity (P=9.5×10^-8^), and Chao (P=0.49), and observed richness (P=0.47), respectively (Table 2). Unsurprisingly, the bacterial diversity and richness also varied by compartment and with watering (Table 2). Fungal alpha diversity was unaffected by the inoculation, nor any of the other experimental factors (Table 2).

**Table 2.** Statistical analysis (Kruskal-Wallis tests and Anova) for the effects of inoculations, plant compartment, and soil water content (SWC) on the bacterial and fungal alpha-diversity.

|  | Bacteria |  |  | Fungi |  |  |
| --- | --- | --- | --- | --- | --- | --- |
|  | Shannon | Chao1 | Observed | Shannon | Chao1 | Observed |
| Compartment (C) | **9.5 x 10<sup>8</sup>\*\*\*** | 0.49 | 0.47 | 0.58 | 0.82 | 0.82 |
| Inoculation (I) | 0.16 | **0.0005 \*\*\*** | **0.0003\*\*\*** | 0.65 | 0.68 | 0.68 |
| SWC | 0.56 | 0.58 | 0.52 | 0.20 | 0.82 | 0.82 |
| Block | 0.99 | 0.66 | 0.65 | 0.43 | 0.48 | 0.48 |

\*\*\*: $P < 0.001$ , \*\*: $0.001 < P < 0.01$ , \*: $0.01 < P < 0.05$ , .: $0.05 < P < 0.10$

We also performed differential abundance analyses (inoculated vs. non-inoculated) at the ASV level, which we visualized using volcano plots (Fig. 3). Each combination of compartment x watering conditions was treated separately. A total of 164 bacterial ASVs showed significant positive responses to inoculation, with 80 ASVs increasing following inoculation under dry conditions and 84 under normal irrigation (Fig. 3A, Supplementary Table S1). The distribution of these positively affected ASVs across plant compartments revealed 71 ASVs (43.3%) in the rhizosphere, 60 ASVs (36.6%) in roots, and 33 ASVs (20.1%) in shoot samples. The lowest P-values were for ASVs belonging to the *Shinella* (P = 5.64 × 10^-17^) and *Rhizobacterium* (P= 2.92 × 10^-14^) genus in the root compartment under dry conditions (Fig. 3, Supplementary Table S1). Conversely, inoculation significantly decreased the relative abundance of 61 bacterial ASVs, with 38 under dry and 23 under normal conditions. The distribution of these negatively affected ASVs revealed 31 ASVs (50.8%) in roots, 22 ASVs (36.1%) in the rhizosphere, and 8 ASVs (13.1%) in shoots. The inoculation negatively affected three ASVs belonging to the *Flavobacterium* genus – two of those showed the most significant decrease among the ASVs affected, with P-values of 4.03 × 10^-6^ and 9.62 × 10^-6^ in roots under normal watering conditions (Fig. 3A, Supplementary Table S2). Three other ASVs from the same genus were positively affected by the inoculation in the rhizosphere and root samples (Supplementary Table S1). Although different isolates of the *Sphingobacterium* genus were in the inoculum, three distinct ASVs from this genus significantly decreased in relative abundance in plant roots and shoots under dry conditions (Supplementary Table S2). Conversely, the only ASV belonging to this genus that was positively affected by inoculation (ASV5) was the one matching the 16S rRNA gene of the inoculated *Sphingobacterium* isolates, which showed a significant increase in the rhizosphere under dry conditions (Supplementary Table S2). Similarly, for the *Paenibacillus* genus, four distinct ASVs decreased after 4 weeks of plant growth, while six different ASVs increased across various plant compartments under both normal and dry conditions (Fig. 3A, Supplementary Tables S1 and S2).

**Figure 3.**
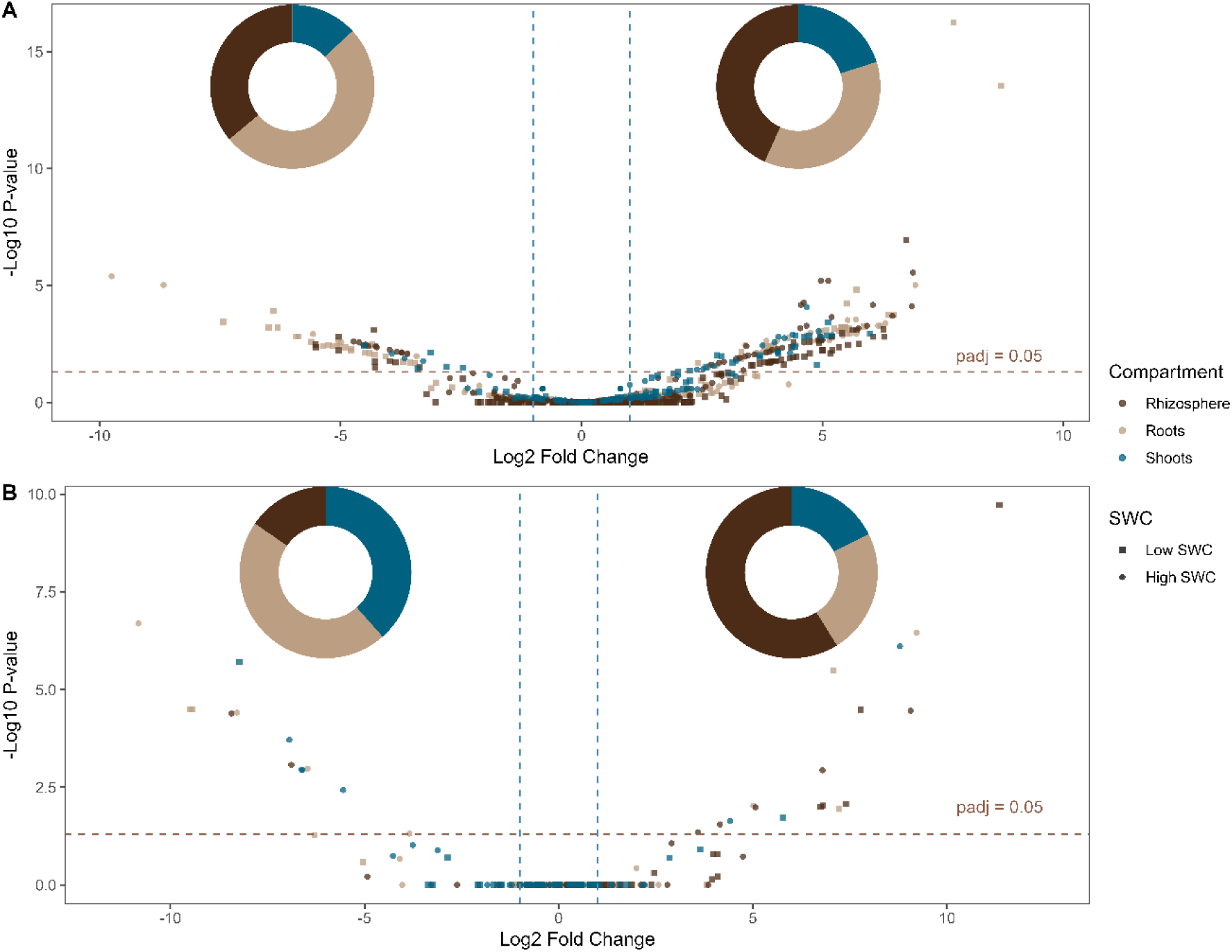
The relative abundance of many microbial ASVs is affected by inoculation. Differentially abundant bacterial (A) and fungal (B) ASVs between inoculated and non-inoculated samples. Differential abundance analyses were performed for each compartment:SWC combinations separately. Donut plots (insert) indicate the proportion for each compartment of positively (right) or negatively (left) differentially abundant ASVs that were found. ASVs with adjusted P-values below 0.05 are listed in Supplementary Tables S1-S4.

Inoculation positively affected 17 fungal ASVs (8 in dry and 9 in normal conditions), from which 10 ASVs were affected in the rhizosphere, 4 ASVs in the roots, and 3 ASVs in the shoots. Conversely, 13 fungal ASVs were negatively affected by the inoculation (3 under dry and 10 under normal conditions), from which 6 ASVs were affected in the roots, 5 ASVs in the shoots, and 2 ASVs in the rhizosphere (Fig. 3B, Supplementary Table S3 and S4). The genus *Gibberella* was the most negatively affected by the inoculation, with 7 different representative ASVs significantly decreasing under both normal and dry conditions across root, shoot, and rhizosphere samples. While the genus *Penicillium* was the most positively affected by the inoculation, with its increase being highly significant under both normal and dry conditions across root, shoot, and rhizosphere samples (Fig. 3B, Supplementary Table S4).

We also tested the effects of irrigation, plant compartment and inoculation on the dominant genera found in the amplicon sequencing datasets (Supplementary Fig. S1 and S2, Supplementary Table S5). We only report here the genera for which the inoculation or its interaction with another factor was significant. The *Klebsiella* was relatively more abundant in the inoculated samples (P = 0.0003), but this also interacted with irrigation (P =0.004) – *Klebsiella* was relatively more abundant in the inoculated leaves under water stress but not so much in the inoculated leaves of well-watered plants (Supplementary Fig. S1, Supplementary Table S5). An interaction between plant compartment and inoculation was found for *Paenibacillus* (p=0.0002). This genus was relatively less abundant in the inoculated samples under drought conditions, but not under normal watering conditions (Supplementary Fig. S1, Supplementary Table S5).

For fungi, the *Zopfiella* genus was affected by inoculation (P =0.036), but in interaction with irrigation (P = 0.01). It was relatively more abundant in the well-watered plants and disappeared in dry plants. Additionally, for the irrigated conditions, this genus was more abundant in non-inoculated plants compared to the inoculated plants (Supplementary Fig. S2, Supplementary Table S5). The *Penicillium* genus was affected by the compartment (P=0.001), with a higher relative abundance in inoculated roots, especially under normal watering conditions (Supplementary Fig. S2, Supplementary Table S5). Under dry conditions, the roots of inoculated plants hosted more *Humicola* (Supplementary Fig. S2) with an interactive effect of inoculation and compartment (P = 0.0003) (Supplementary Fig. S2, Supplementary Table S5). Similarly, the *Epicoccum* genus, was relatively more abundant in the inoculated rhizospheres under non-irrigated conditions compared to irrigated conditions (Supplementary Fig. S2, Supplementary Table S5) indicating an interactive effect between inoculation and plant compartments (P = 0.003).

### Genomic analysis of the inoculated microbes

We sequenced the genomes of the inoculated microorganisms. The microorganisms were identified using the average nucleotide identity (ANI) against sequenced isolates available in GenBank (Table 3). We also looked in the whole genomes for their 16S rRNA genes or ITS region and compared them to the amplicon dataset to match each inoculated microorganism to an ASV. Among the 25 inoculated strains, 5 bacteria and 2 fungi had no match to ASVs, whereas 18 bacteria isolates had a 100% similarity with a unique ASV (Table 3). Thereby, we could determine which ASV from the inoculum potentially persisted in the plant environment and which were not detectable anymore at the time of sampling. Some of the inoculated microorganisms matched the same ASV because they were closely related and had highly similar 16S rRNA genes. They also matched the same published genome based on the ANI analysis, potentially indicating different strains from the same species (Table 3).

**Table 3.**
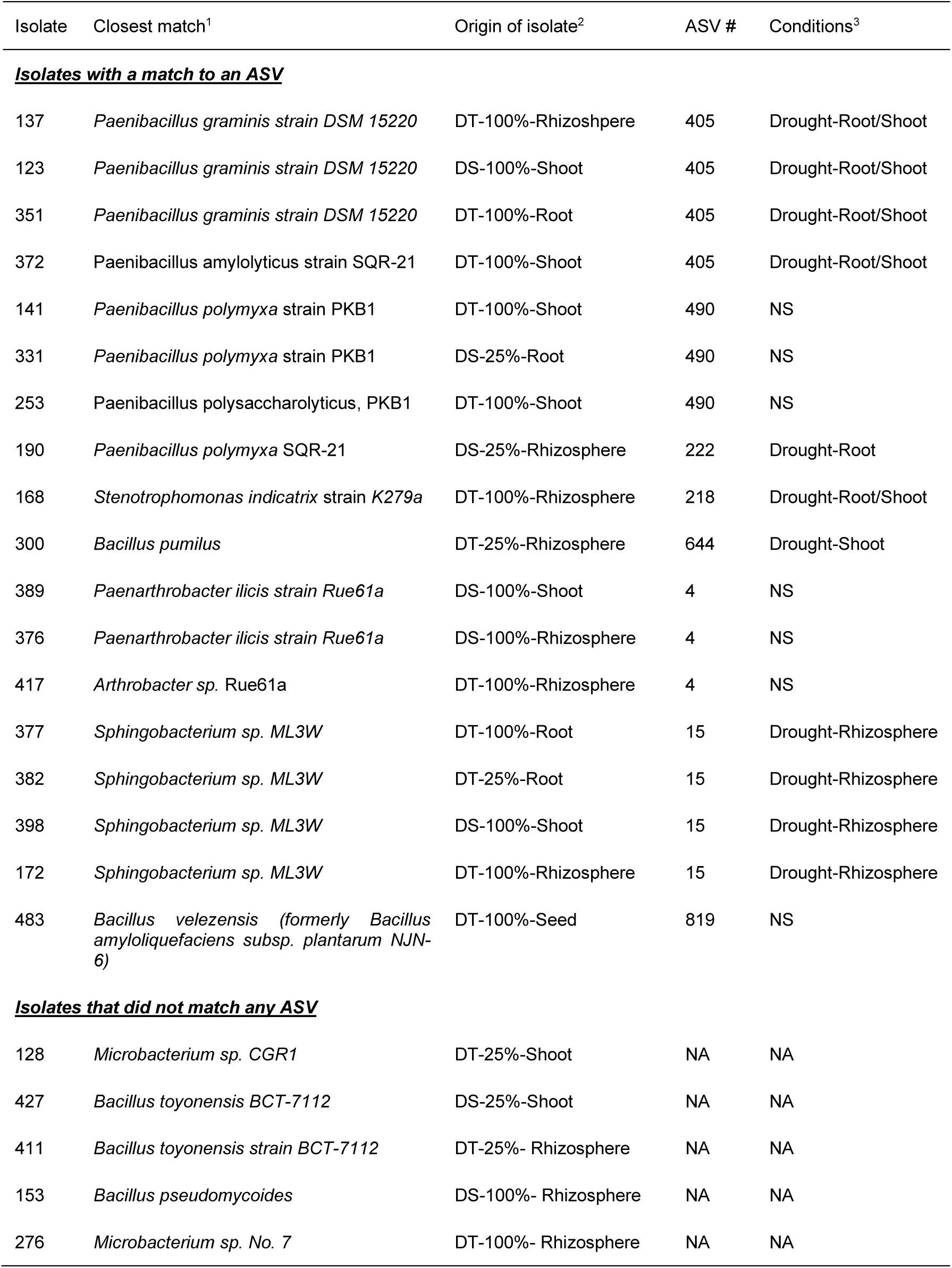
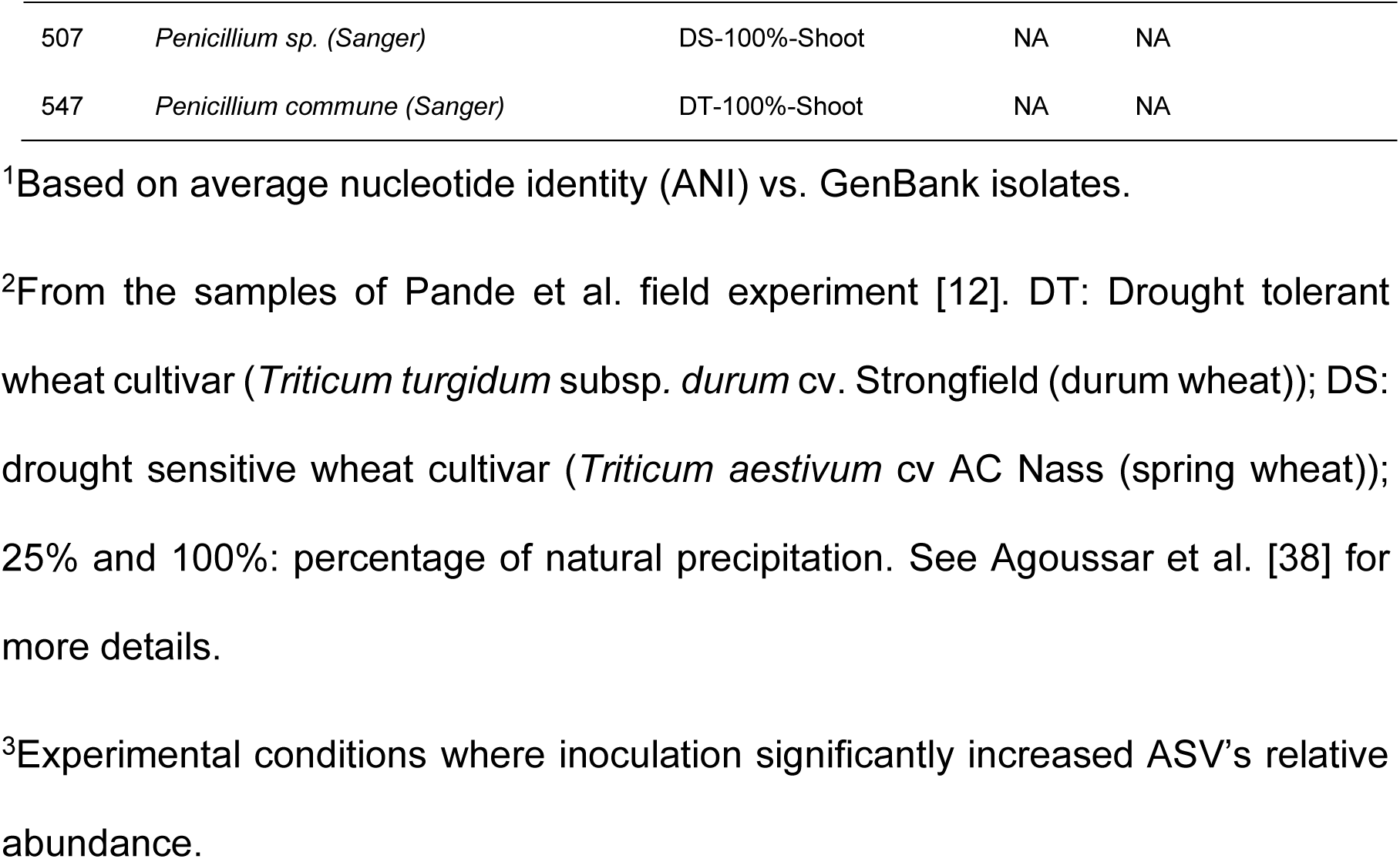
Bacteria isolates used to prepare the “Screening” inoculum, their origin, the corresponding ASV having 100% similarity with their 16S rRNA gene, and the conditions.

None of the ASVs matching the inoculated isolates decreased in relative abundance following inoculation. Across various compartments, several of the ASVs corresponding to the inoculated isolates showed significant increase following inoculation, and this was only seen under low water availability (Table 3). For instance, ASV405, matching different isolates identified as *Paenibacillus graminis* (isolates 123, 137, 351), showed a significant increase in relative abundance in both roots and shoots under water deficiency. Other ASVs, such as ASV218 and ASV644, matching the isolates *Stenotrophomonas indicatrix* and *Bacillus pumilus* respectively, showed a significant increase in relative abundance in shoots under low water conditions. Similarly, ASV15, which matched different *Sphingobacterium* isolates, significantly increased in relative abundance in the rhizosphere, also in the low water treatment.

Based on their detection in the plant environment at the end of the experiment, the isolates used to prepare the inoculum were separated into two groups. The first group consisted in microbes that had no match with the ASVs retrieved from the amplicon sequencing (“non-persisters”) and the second group consisted in microbes that had a match with an ASV at 100% (“persisters”). The microbes from the first group probably did not colonize the wheat environment to grow in sufficient numbers to be detectable, whereas the microbes in the second group potentially persisted in the wheat environment to reach numbers above the detection level. This comes with the caveat that the “non-persisters” might have been present, but below the detection limit of the method, whereas the “persisters” might have matched the 16S rRNA gene of a closely related ASV from the environment. With that limitation in mind, we compared the genomes of the two groups to identify potential key genomic factors implicated in persistence, but also in the larger plant biomass under water stress. As compared to the non-persisters, the potential persisters had larger genomes (P=0.012) that contained more genes (P=0.094), and more of these genes were part of orthogroups (P=0.055) (Fig. 4).

**Figure 4.**
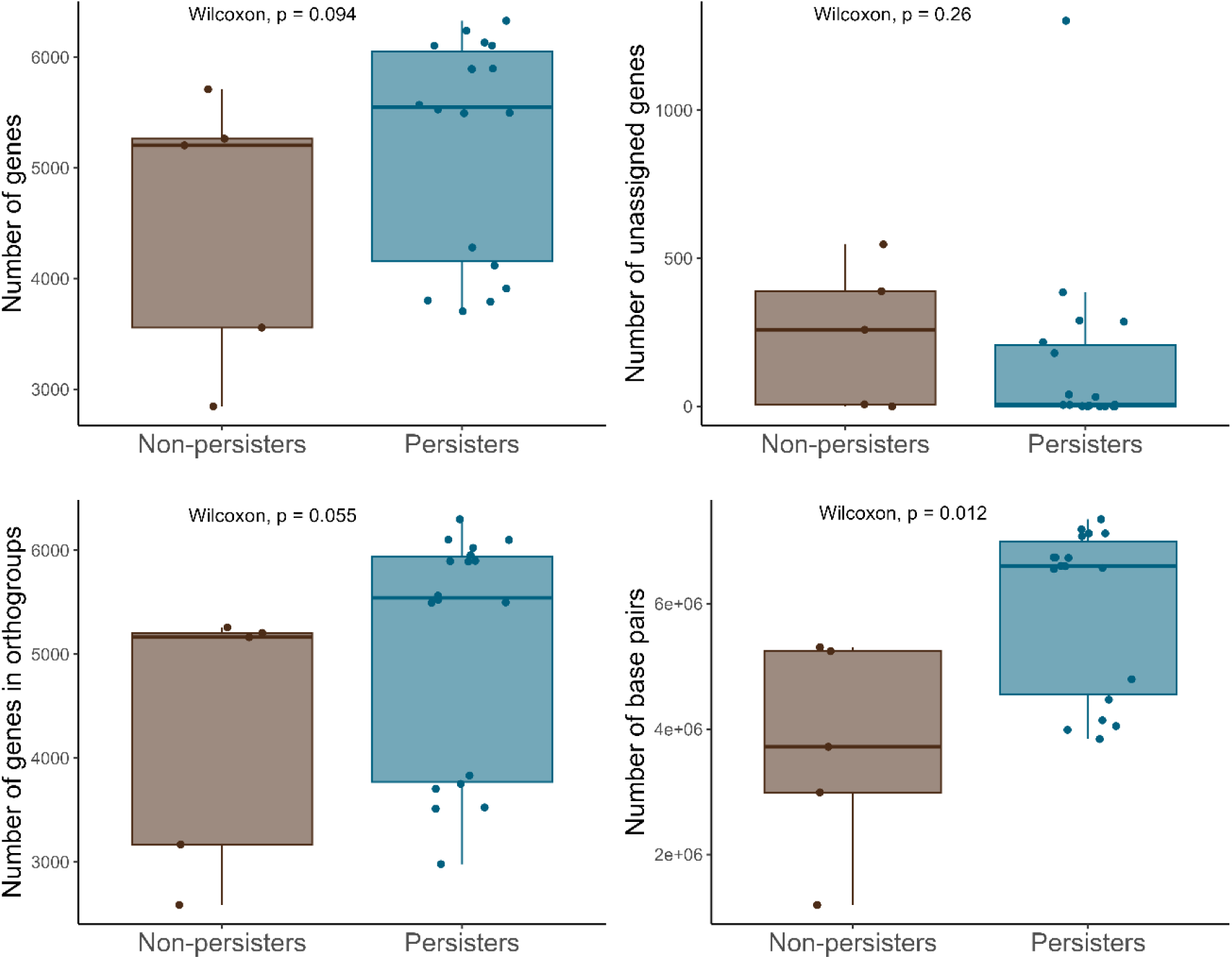
Inoculated microorganisms that persisted in the wheat environment share genomic characteristics. Genomic characteristics of the 23 inoculated bacteria, grouped by their persistence in the wheat environment (defined as finding a perfect match to the 16S rRNA gene of an ASV).

We identified 159 genes common to the 23 bacterial strains. We also looked at the genes commonly found in the “persisters” but totally absent in the “non-persisters”. Some were shared among more than three quarters of potential “persisters” (more than 14 out of the 18 isolates) and absent in the “non-persisters” (Fig. 5). We also investigated differences between “persisters” and “non-persisters” in the presence of secondary metabolites biosynthetic gene clusters using the antiSMASH tool. We found out that some secondary metabolites gene cluster predicted for the “persisters” with similarity scores exceeds 75%, were absent among the “non-persisters” (Fig. 6). The “persisters” shared several clusters associated with antimicrobial activity, such as colistin, bacillopaline, lichenysin, macrolactin, polymyxin, and thermoactinomide A. The “non-persisters” also exhibited clusters linked to siderophore production, such as bacillibactin and petrobactin. Interestingly, the isolate *Sphingobacterium sp.* 398 did not contain any known secondary metabolite gene cluster with a similarity score above 75%, despite being categorized as a “persister”. In contrast, isolate *Bacillus velezensis* 483 contained seven known secondary metabolite gene clusters (bacillaene, bacillibactin, bacilysin, difficidin, fengycin, macrolactin, and surfactin) with similarity score above 75%.

**Figure 5.**
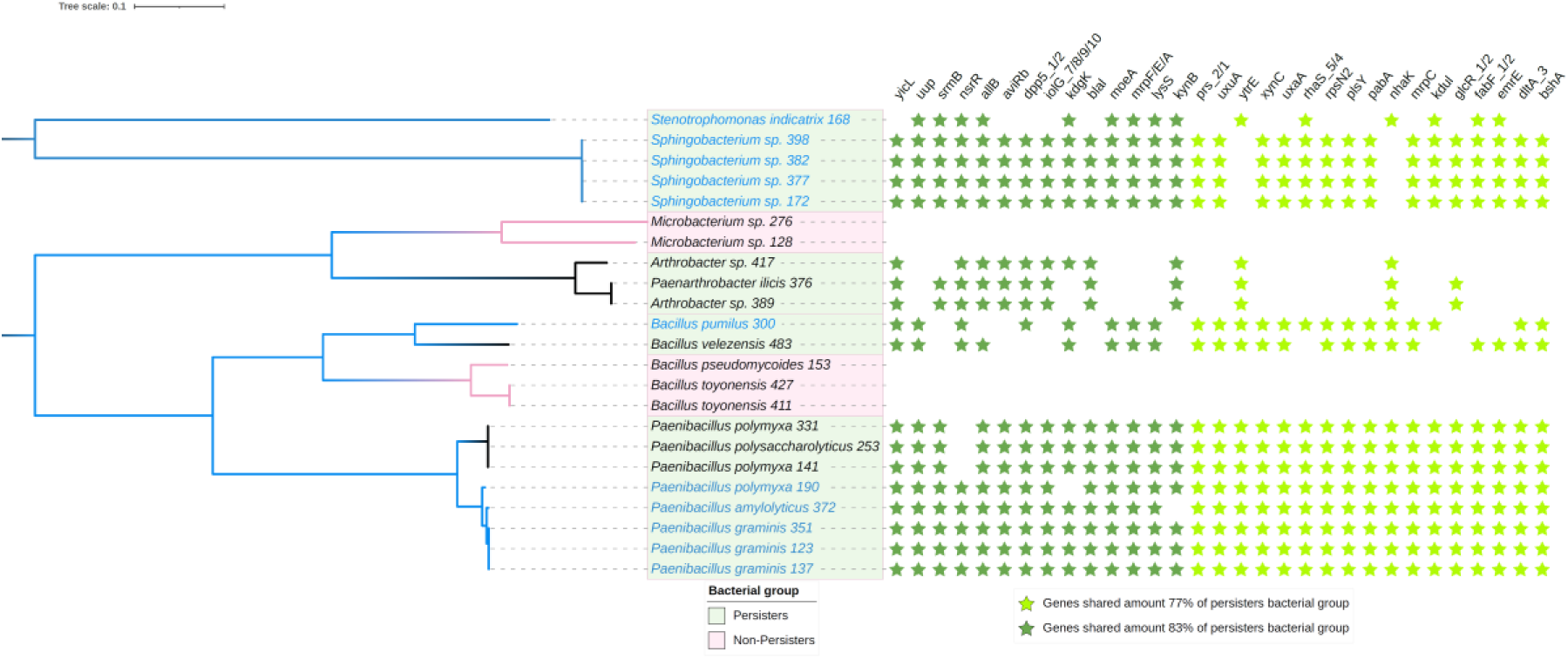
Inoculated microorganisms that persisted in the wheat environment share genes that are absent in the microorganisms that did not persist. Phylogenomic tree illustrating genetic relationships among the 23 bacterial isolates based on a comparative genomic analysis of amino acid annotations for their genomes. Branch lengths represent the genetic distances between isolates. Bacterial isolates highlighted in green persisted in the wheat environment (defined as finding a perfect match to the 16S rRNA gene of an ASV), while those in pink did not. Among the persisters, isolates in blue letters are those for which their corresponding ASV was significantly more abundant in the 15% SWHC treatment compared to the 50% SWHC treatment. Key genes shared by 14 out of 18 persisters (77%) are indicated with light green asterisks, and those shared by 15 out of 18 persisters (83%) are marked with dark green asterisks. In both cases the genes are absent in the non-persisters.

**Figure 6.**
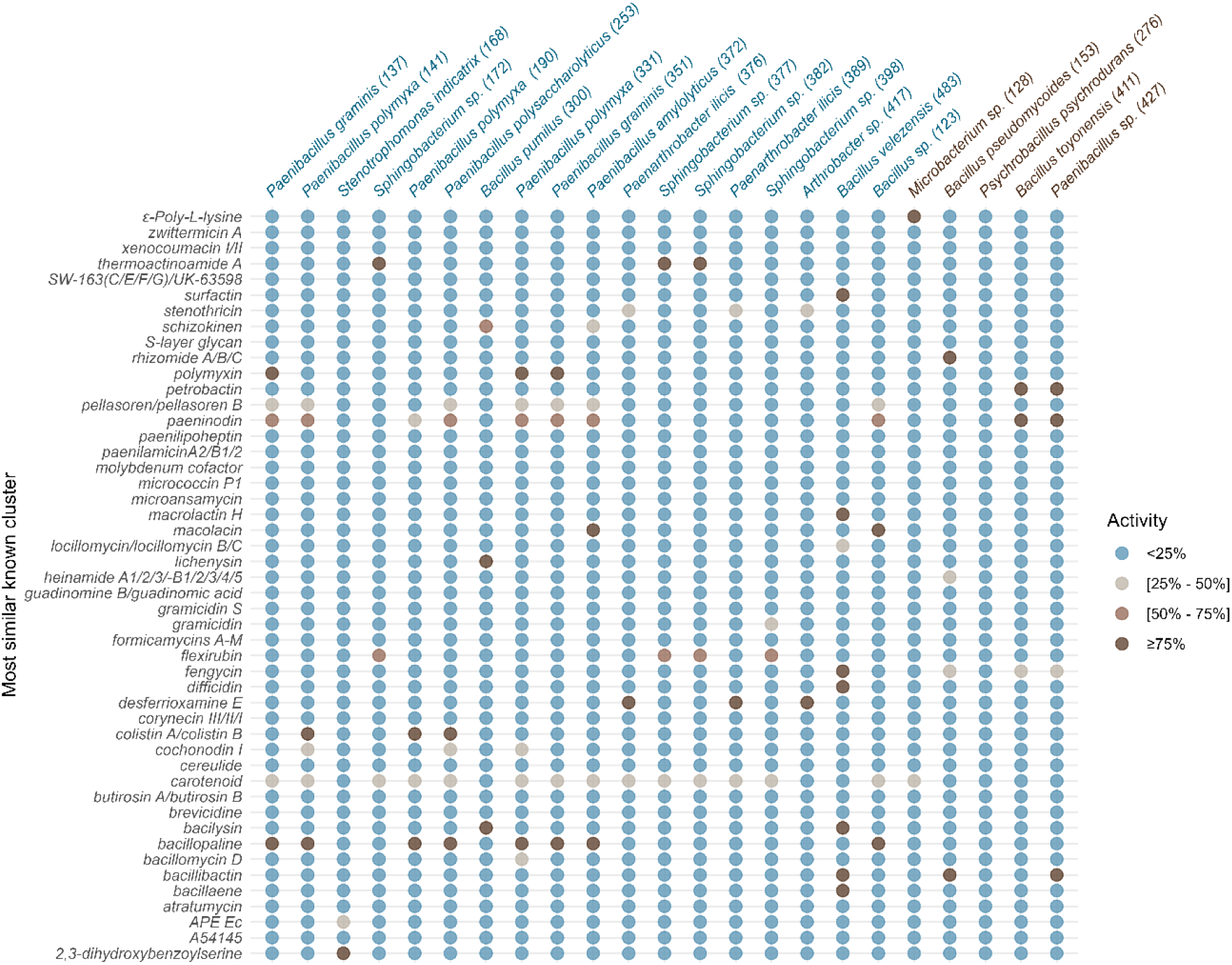
Persisters’ genomes generally contained more secondary metabolites genes than the ones of non-persisters. Most similar known secondary metabolite gene clusters in the genomes of persister (blue, defined as finding a perfect match to the 16S rRNA gene of an ASV) and non-persister (brown).

## Discussion

Plant and soil associated microbes have a key role in helping crops adapt to abiotic stresses, such as drought [48]. One way to maximize yields under the current climatic emergency would be to manipulate the plant microbiome. Although the field is still in its infancy, a general ecological framework was suggested [3], which included migration (addition of new microorganisms) and selection (changes in the resident microbial community), two ecological mechanisms that could explain the effects of inoculants. Here, we compare these two mechanisms. We found that, for the inoculum that had worked best, both migration and selection were likely. Many genomic features of the inoculated organisms that had potentially persisted, as well as the shifts observed in the plant microbiota could explain the increased plant growth under water stress.

First, we compared inoculants that were created using different approaches. Many different methods are available to create inoculants [39], but they are rarely tested side-by-side. In a previous study, we compared microbes extracted from a soil with a water stress history or not (a naturally “evolved” inoculum), that were inoculated to wheat plant under water stress [9]. This resulted in no improvement in plant growth or water content and only modest shifts in the fungal communities in the rhizosphere [9]. We had also applied this approach in the context of bioremediation of hydrocarbon petroleum, with similar lack of improvement in plant growth and hydrocarbon degradation rates using an “evolved” inoculum [49,50]. In fact, for both studies, the control inoculum that was not “evolved” was more efficient to promote plant growth or degrade hydrocarbons [49,50]. We suggested that a highly diversified, unselected inoculum was more prone to contain the optimal microbes for the process of interest [49,50]. Mismatches between the successional stages of the selected communities and of the inoculated ecosystem were also suspected to create problems. We also found here that the inoculum created by “evolving” soil under dry or wet conditions had little effect on wheat growth under water stress. However, in contrast to our previous studies, we did not include an unselected, highly diversified inoculum, but included instead a diversified inoculum that was created from 25 isolates that were able to grow under high osmotic pressure. This targeted approach – the “screening” inoculum – was the only one that led to improved wheat biomass under water stress.

The inoculum developed from isolates that grew under high osmotic pressure was able to promote plant growth, by increasing wheat aboveground fresh biomass under water stress, but not the dry biomass nor the root biomass. This indicates that the “Screening” inoculum probably helped wheat to retain water in its leaves. This was also evident when looking at the plant morphology, with the inoculated plants showing an improved turgor under water stress. The inoculum could have therefore stimulated stomatal closure or helped plant accumulation of osmolytes, two mechanisms that are compatible with our observations [51,52]. Simply promoting plant growth is not an ideal mechanism to help plants survive drought, because plant communities with more biomass are more susceptible to drought [53]. Our screening method, using high osmolarity growth media, selected for microbes that were highly efficient in producing osmolytes. Microbial endophytes and rhizobacteria can increase plant osmolyte concentration [31–33], and can also exsude osmolytes in the plant environment [30,34]. We also recently showed that microbial osmolytes-related transcripts were more abundant in the wheat rhizosphere when soil water content decreased [12], and that intermittent water stress selected for rhizosphere microorganisms that were enriched in osmolyte producing genes [13].

Among the bacterial strains inoculated that were represented among the ASVs (the “persisters”), twelve out of seventeen were Gram-positive bacteria. Gram-positive bacteria produce osmolytes constitutively whereas Gram-negative bacteria produce them as a drought-induced response [54]. This constitutive production of osmolyte together with a thick peptidoglycan cell wall in Gram-positive bacteria allows them to remain active under low water availability, in contrast to bacteria that avoid drought by dormancy or sporulation [55]. Only active bacteria can protect the plant from water stress, suggesting that bacteria that have a drought resistance rather than a drought avoidance strategy would be better inoculants.

We had screened a collection of 550 isolates for growth on a hyperosmolar media containing 30% of PEG [38], which resulted in the 25 isolates (23 bacteria and 2 fungi) used here. Genomic analysis of the 23 bacterial isolated used in the “screening” inoculum highlighted several genes that could be linked to the capacity of the isolate to live under low water availability. For instance, they all contained the “*BetI”* gene, responsible of choline-responsive regulator that controls the synthesis of glycine betaine, enabling osmoadaptation under hyperosmotic stress conditions [56–58]. The “*obg”* gene was also shared by the 23 isolates, and plays a key role in the bacterial stress response by helping activate the sigma B (σ˄B) protein in response to environmental stress [59]. Our genomic analysis highlighted several candidate genes that could be further examined to confirm their causal role in the capacity of our isolates to live under low water availability.

Having traits related to growth under water stress is not the only thing needed for microorganisms to form a successful inoculum for helping crops resist to water stress. The microorganisms also need to establish themselves and hopefully thrive in the plant environment when they are applied. Isolates that are good candidates for improving the plant phenotype are not often screened for this trait. So, we used the differences in potential persistence among our multi-species inoculum to try to understand which genomic factors were important for successful establishment. The genomes of the “non-persisters” clustered together, and they missed several of the genes that were widely shared among the “persisters”. Many “persisters” had the genetic potential to produce different antibiotics, such as *colistin, bacillopaline, lichenysin, macolacin, polymyxin,* and *thermoactinomide A*. and other secondary metabolites, such as siderophores. The presence of these genes in the genome of the “persisters” is well aligned with the traits that are required to colonize the plant environment. For instance, the capacity of *Paenibacillus polymyxa* to promote the growth of plants was partly linked to its capability to produce polymyxin [60]. It makes sense that the potentially persistent isolates shared these traits in their genomes. These genes and the resulting traits could be targeted when screening new isolates, to increase the chance that they might establish in the plant environment.

On top of their direct effects on plants, inoculants can modify the resident soil and plant microbial communities [15,16]. For instance, *Bacillus* can shape the microbial community in the rhizosphere [61], whereas *Paenibacillus polymyxa* produces antibiotics [62], which could also modulate the microbial community. In our study, this could explain indirectly the effect of our “Screening” inoculum, as inoculation affected the bacterial and fungal communities. For example, the relative abundance of the *Shinella* genus was positively affected by the inoculation, most especially in wheat roots under water stress. Bacteria from this genus significantly enhanced duckweed biomass and root development [63] and were enriched in the rhizoplane of wheat [64]. Some strains of *Klebsiella* can accumulate osmolytes such as glycine betaine, trehalose and proline in response to drought stress [30], and, here, this genus was relatively more abundant in leaves following inoculation. Different genera from the Rhizobiaceae family such as *Allorhizobium, Neorhizobium, Pararhizobium,* and *Rhizobium* showed a significant increase in their relative abundance in root samples under dry conditions following the inoculation. These genera are known to increase plant osmolyte concentration [31–33] and to play a crucial role in supporting plant growth in nutrient-poor and drought-prone environments [65]. Additionally, among the nine fungal ASVs that were negatively affected by the inoculation, five of them belonged to the *Gibberella* genus, a known wheat pathogen [66,67]. Many of the inoculated bacteria had the genetic potential to produce secondary metabolites that could affect fungi. For instance, *Paenibacillus polymyxa* can protect cereals against *Fusarium* head blight caused by *Fusarium culmorum* [68].

The genera listed above were not part of our inoculum and were therefore amplified from environmental strains following inoculation. More intriguing were the shifts observed in ASVs from bacterial genera that were represented in our inoculum. The ASVs matching our isolates never decreased in relative abundance following inoculation, but ASVs from the same genus were sometime negatively affected. This was the case for several ASVs from the *Paenibacillus, Sphingobacterium, Bacillus*, *Stenotrophomonas,* and *Penicillium* genera, which were well represented in our inoculum. Since these bacterial genera can enhance drought stress resistance in plants [60,69–74], and *Penicillium* can help plant accumulate osmolytes such as proline under drought [75–77], these negative effects could have reduced the positive effects of the inoculation. It would be interesting to further understand the role of niche and taxonomical overlap between the inoculated and native microorganisms on the persistence and efficiency of the inoculated microorganisms, and on its effects on the native community.

We found that, among the four approaches tested, only the inoculum made of a mixture isolates able to grow at high osmolarity and promote plant growth successfully enhanced wheat aboveground fresh weight. Not all strains of the inoculum potentially persisted in the plant environment, and this persistence could be linked to genomic features. The question remains as to whether the inoculated strains acted directly on the plant, or indirectly through shifts in the microbial communities. Our data supports both mechanisms, and probably the effect on plant fresh biomass was a result of a combination of the two mechanisms. Microbiome engineering approaches that combine more than one mechanism of action are more likely to be successful [2,3], which could provide direly needed tools to adapt crops to the ongoing climatic emergency.

## Acknowledgments

We are grateful to Liliana Quiza and Pranav Pande for help with sampling the experiment.

## Data availability statement

The genomes of the isolates are available under the NCBI BioProject accession PRJNA884320. The amplicon datasets are available under the NCBI BioProject accession PRJNA1214753.

---

**Figure S1.**
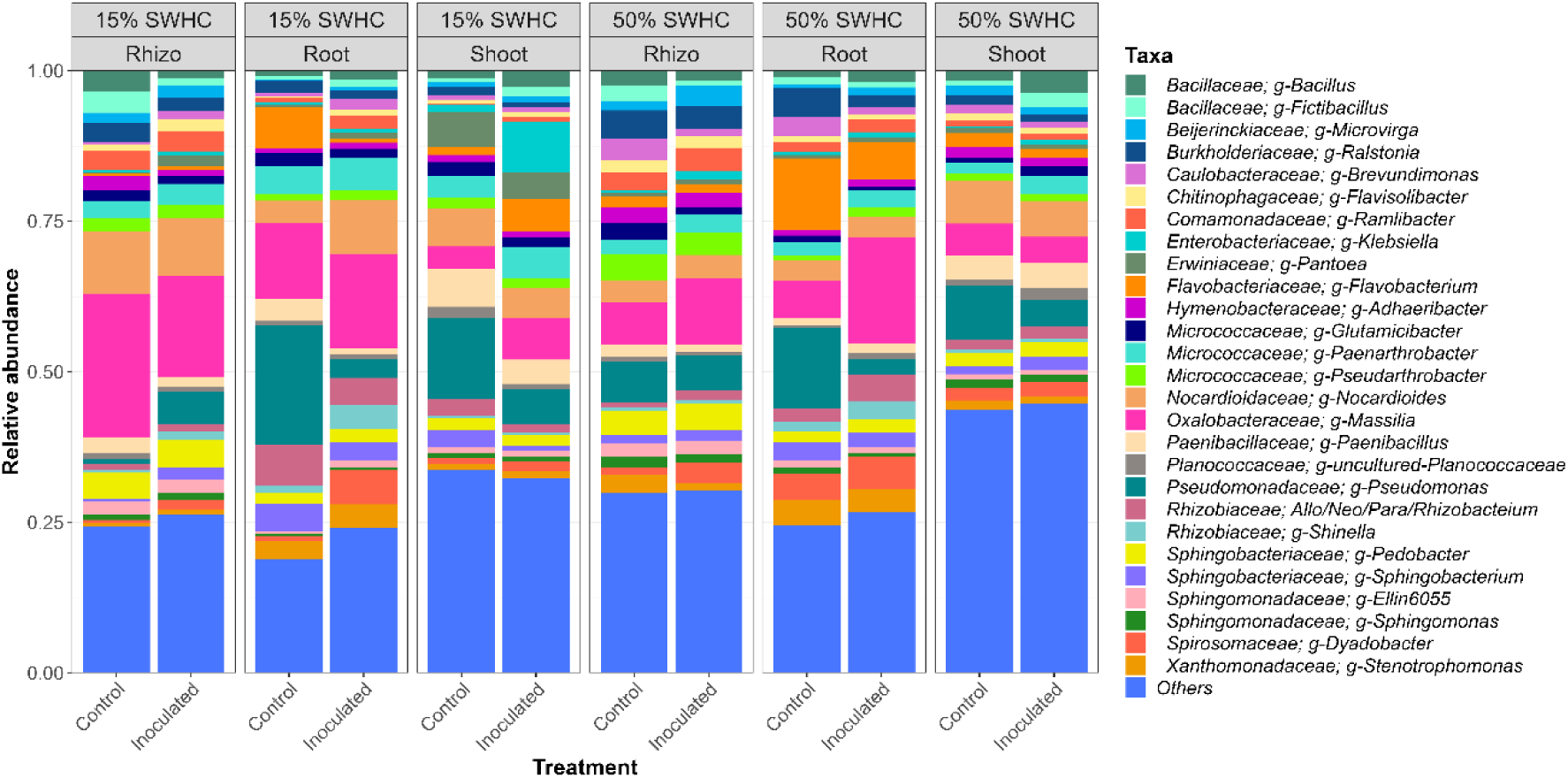
Relative abundance of the dominant bacterial genera (on average, more than 1% of the 16S rRNA gene reads) retrieved from shoot, root and rhizosphere samples from wheat inoculated or not with and growing under two different soil water content (15% and 50% SWHC).

**Figure S2.**
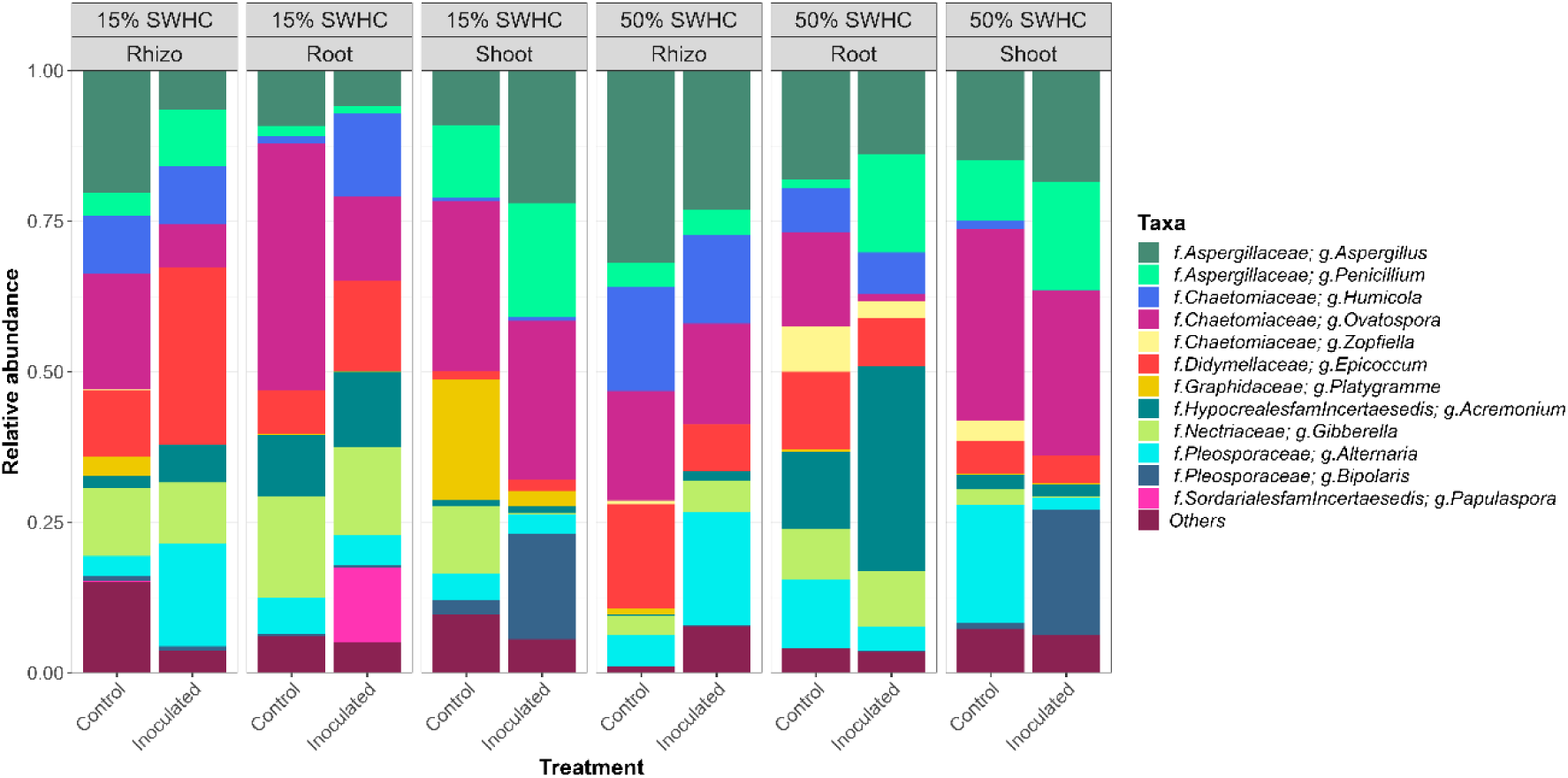
Relative abundance of the dominant fungal genera (on average, more than 1% of the ITS reads) retrieved from shoot, root and rhizosphere samples from wheat inoculated or not with and growing under two different soil water content (15% and 50% SWHC).

## Supplemental material

**Table S1:**
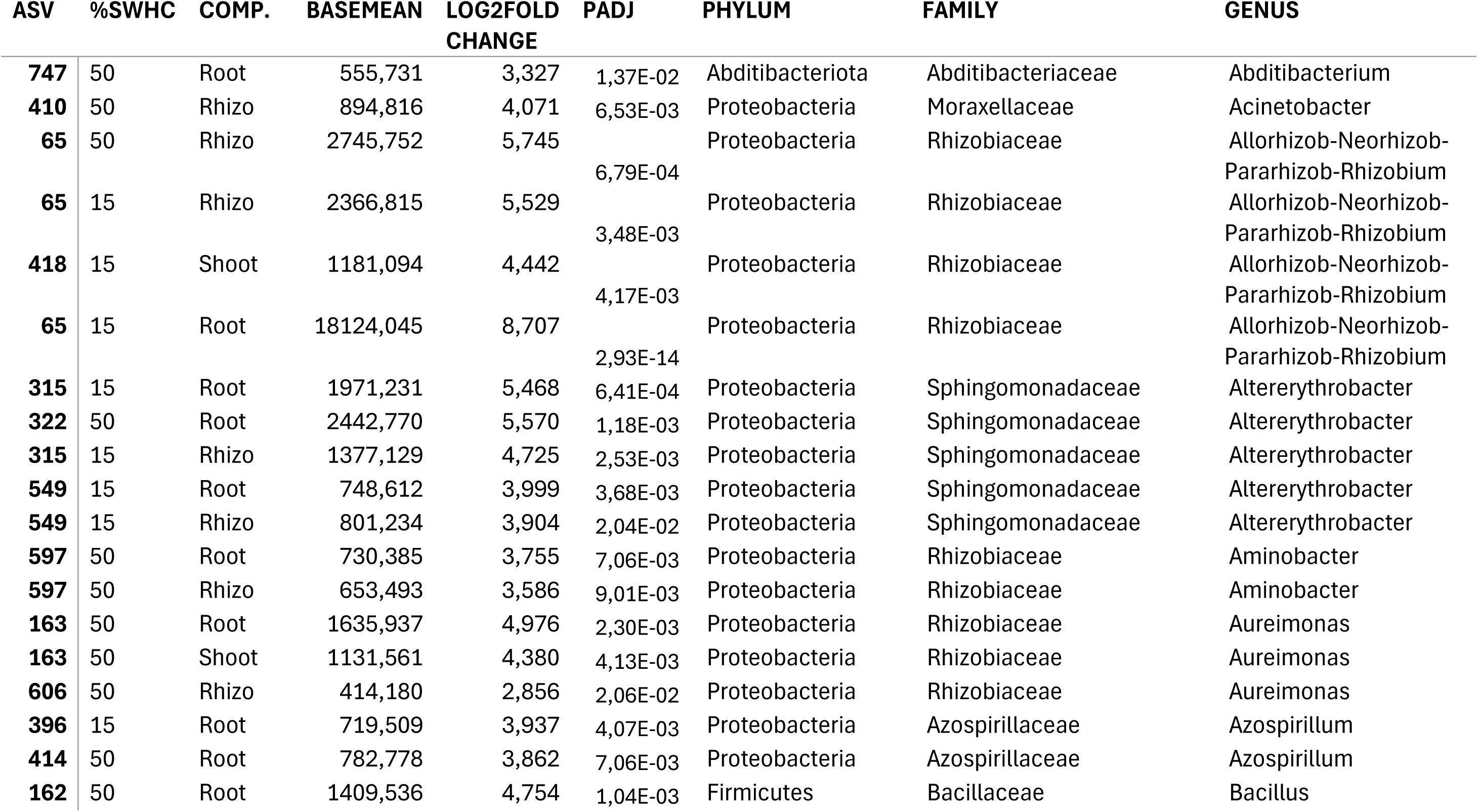
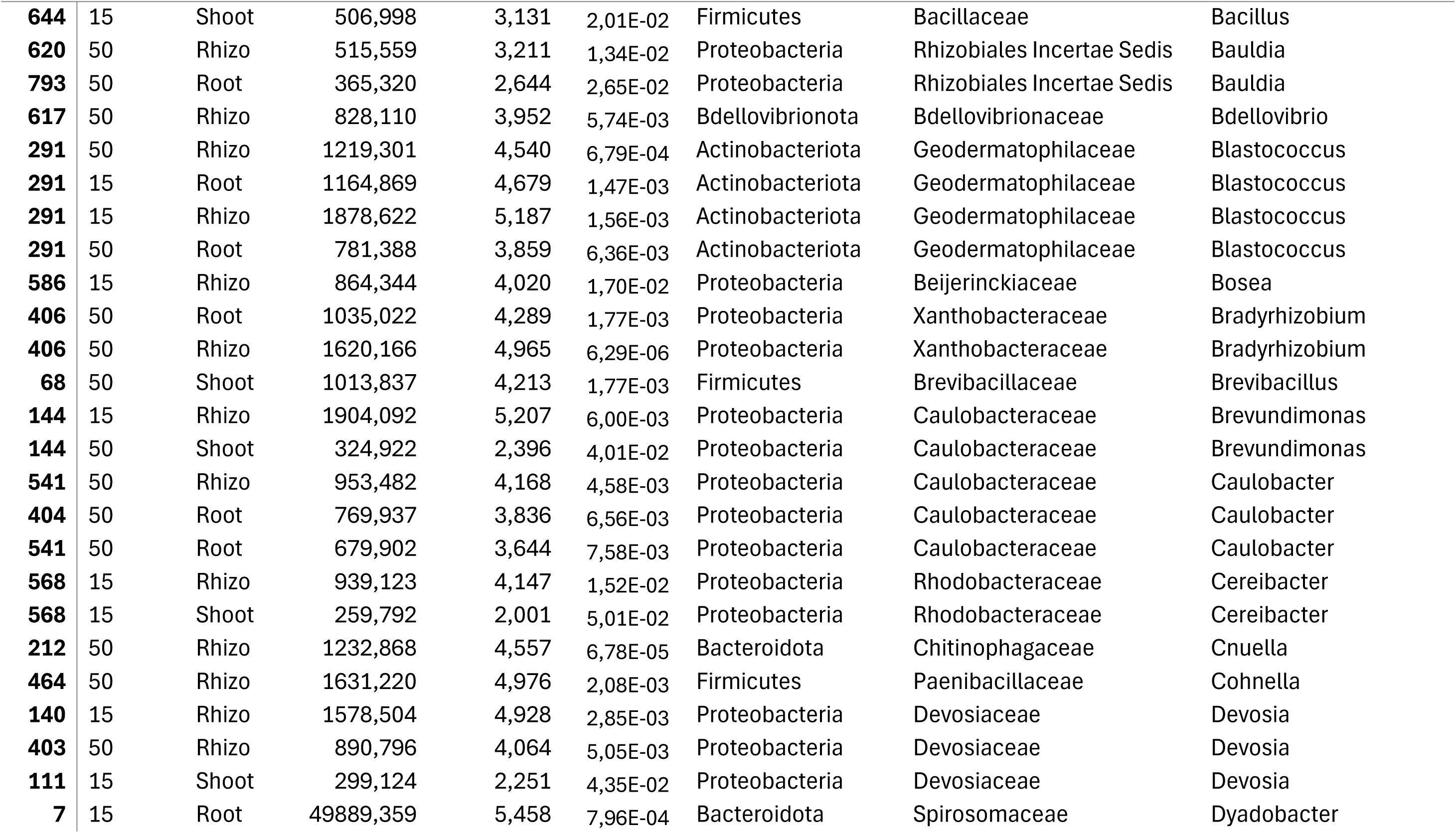
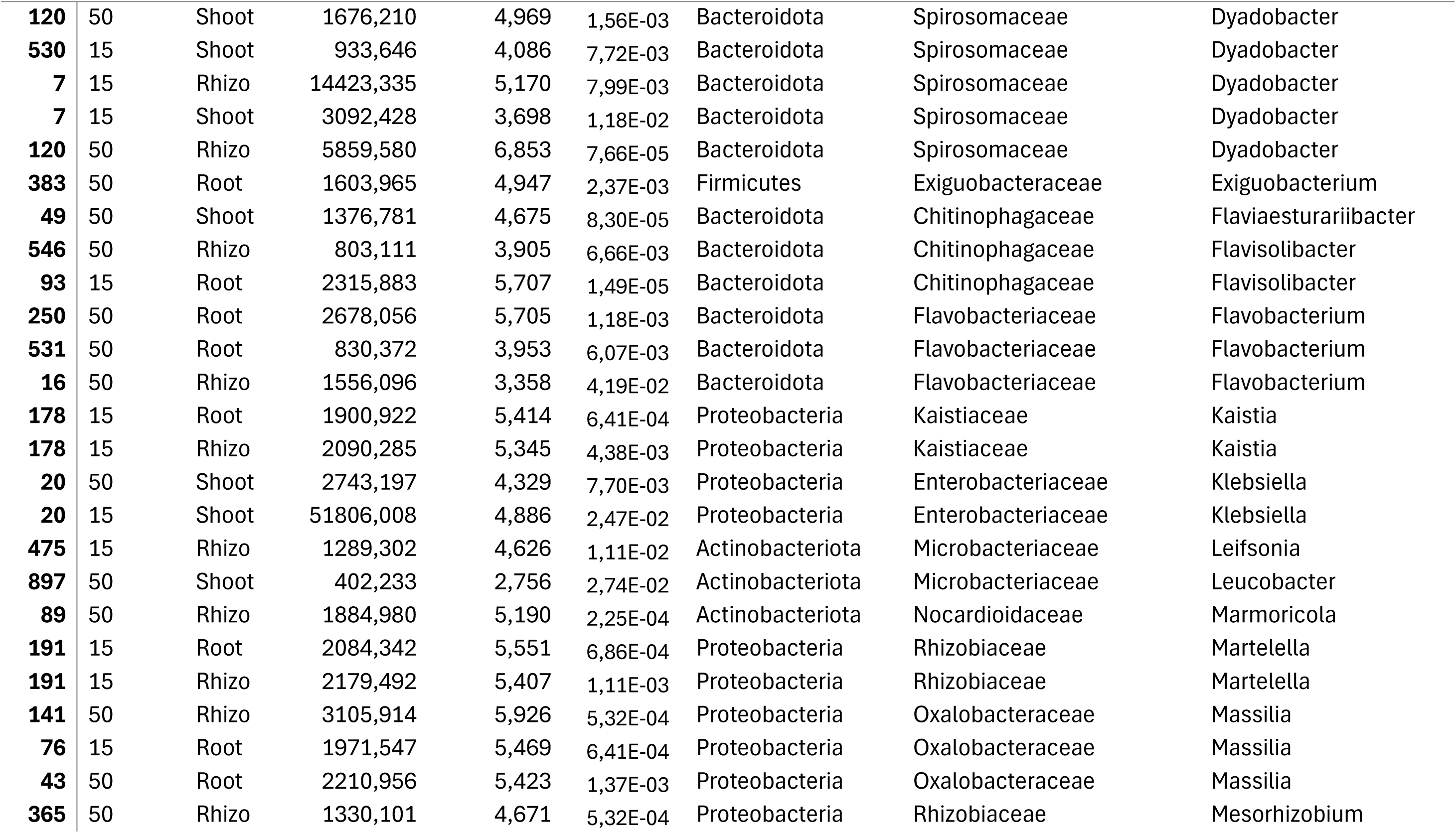
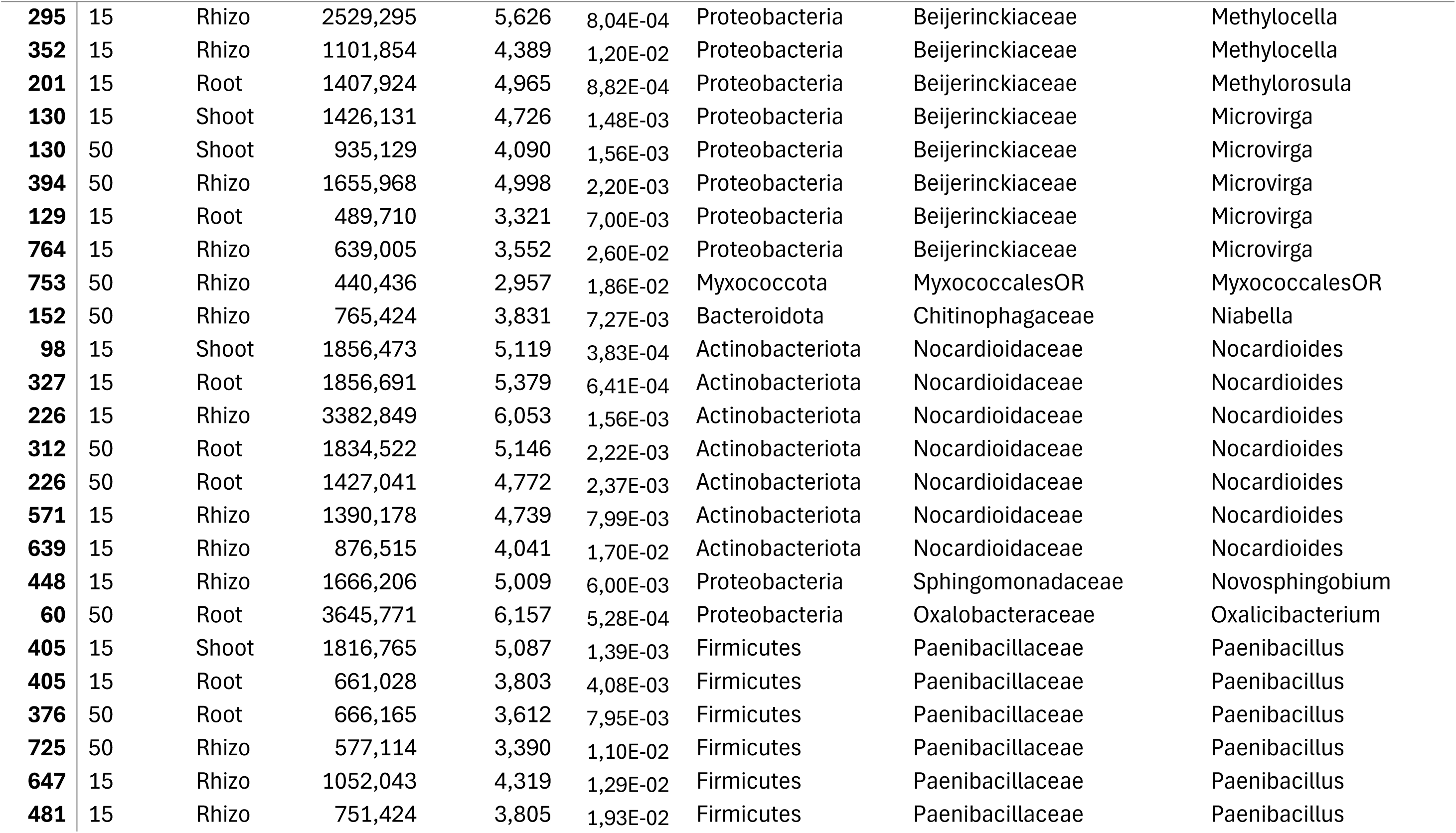
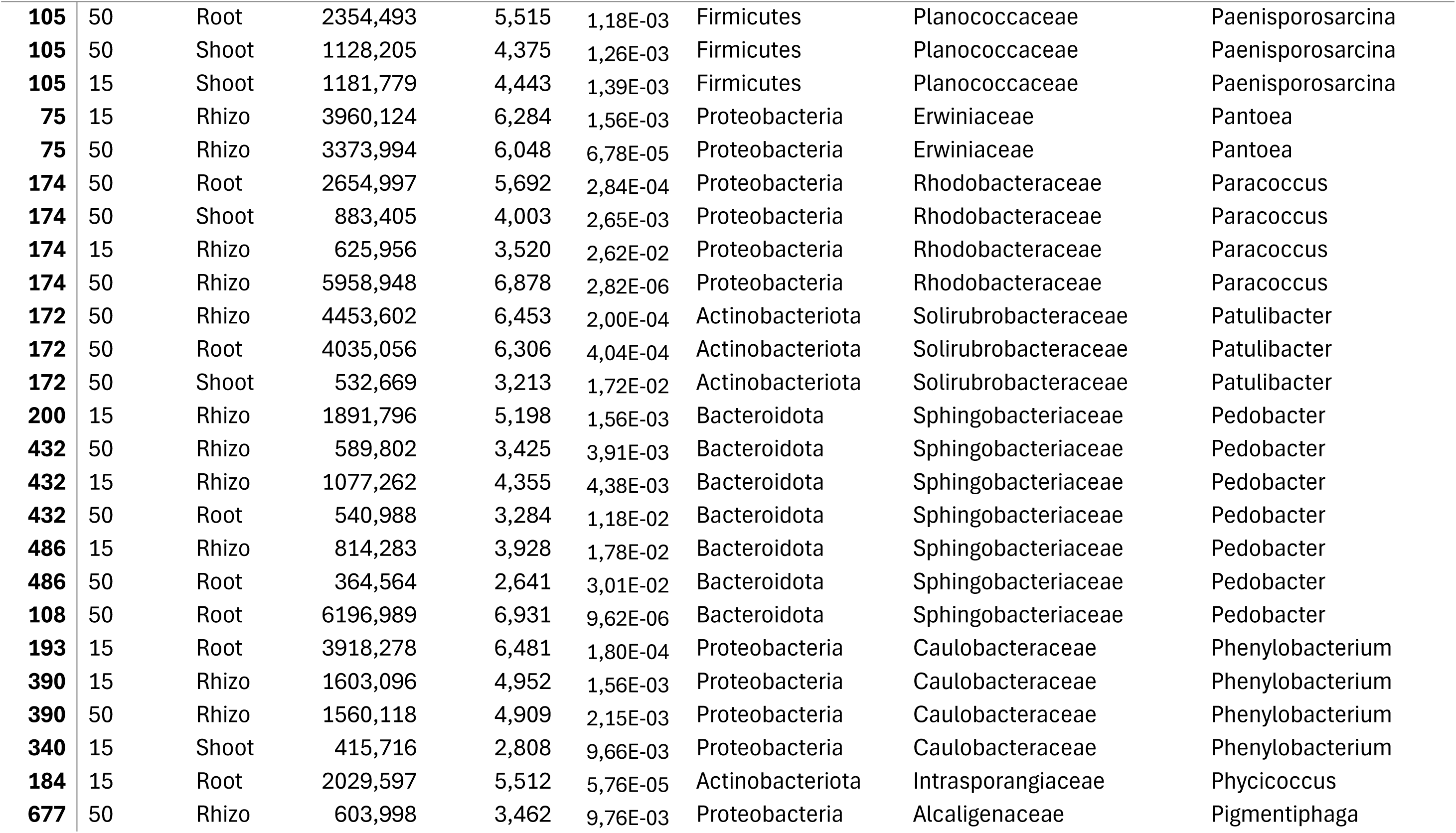
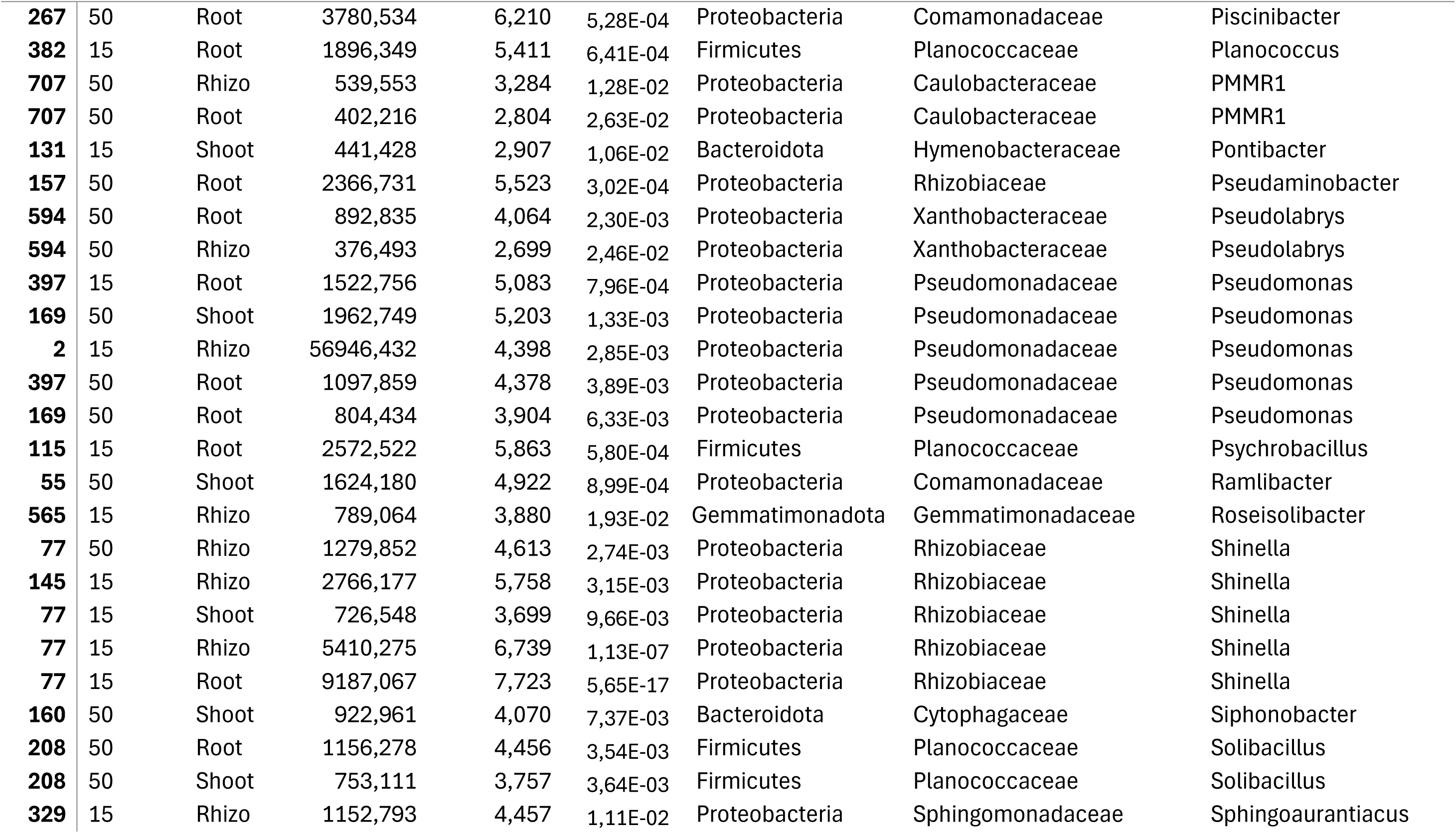
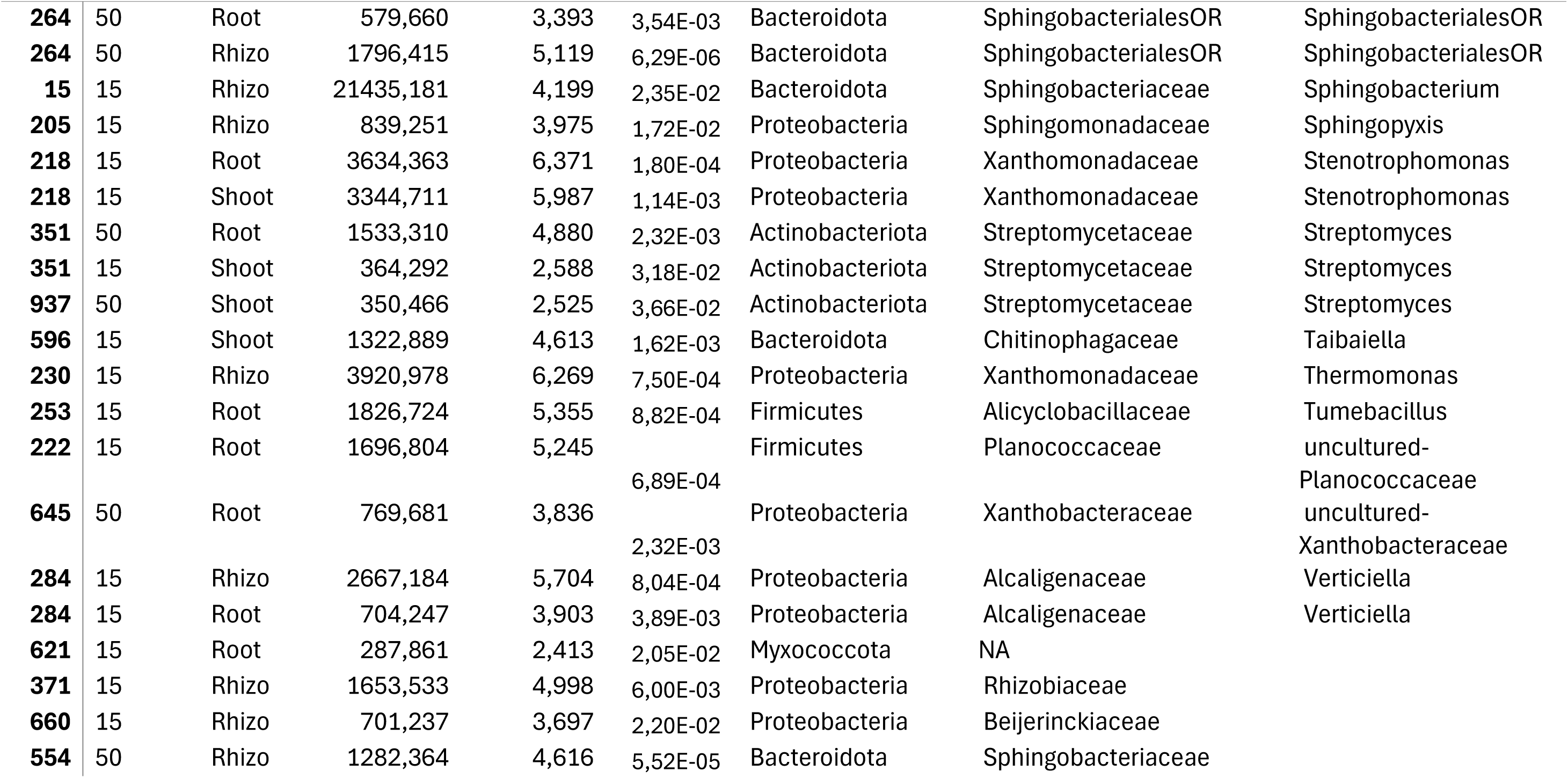
Bacterial ASVs positively affected by the inoculation.

**Table S2:**
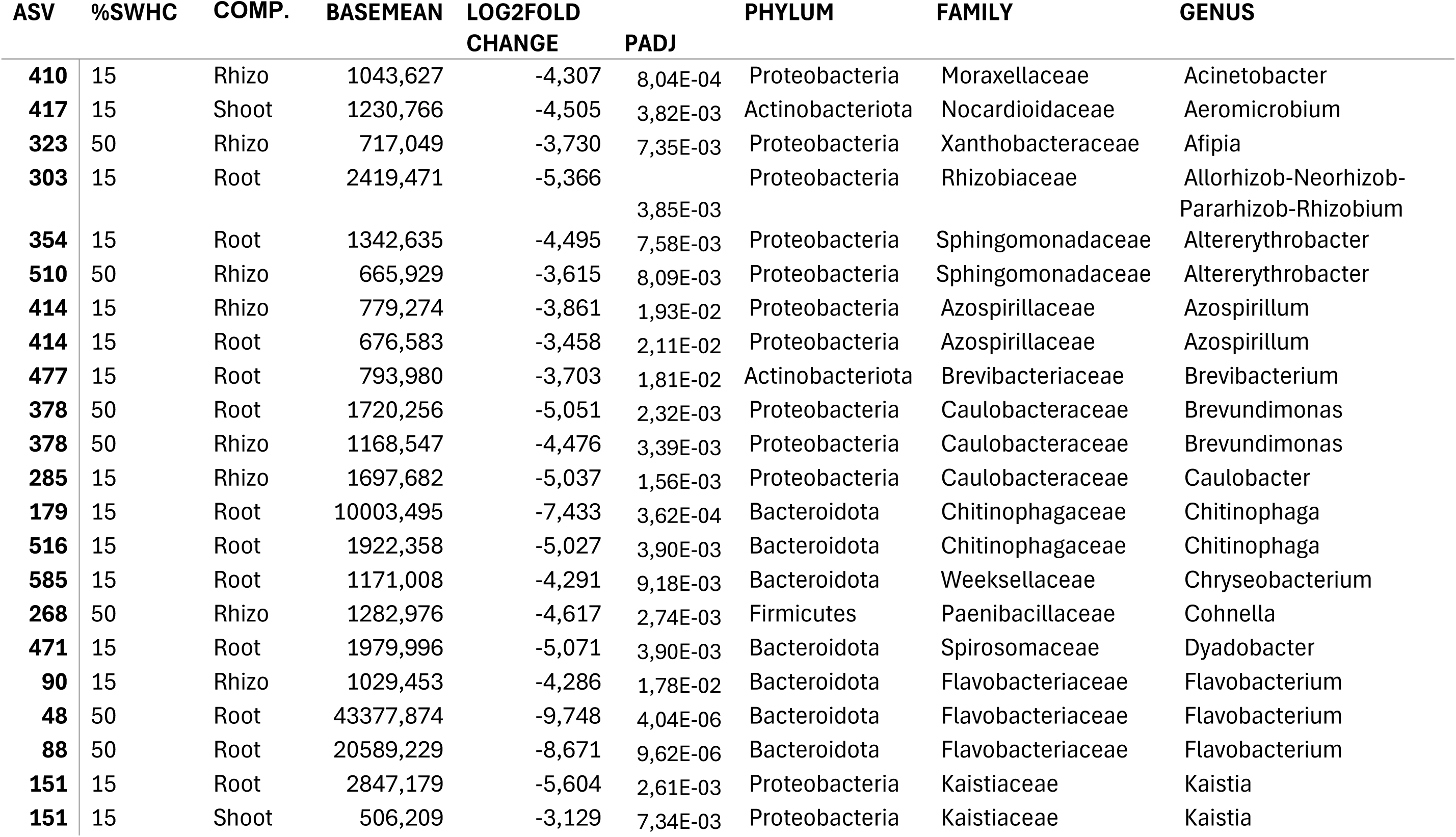
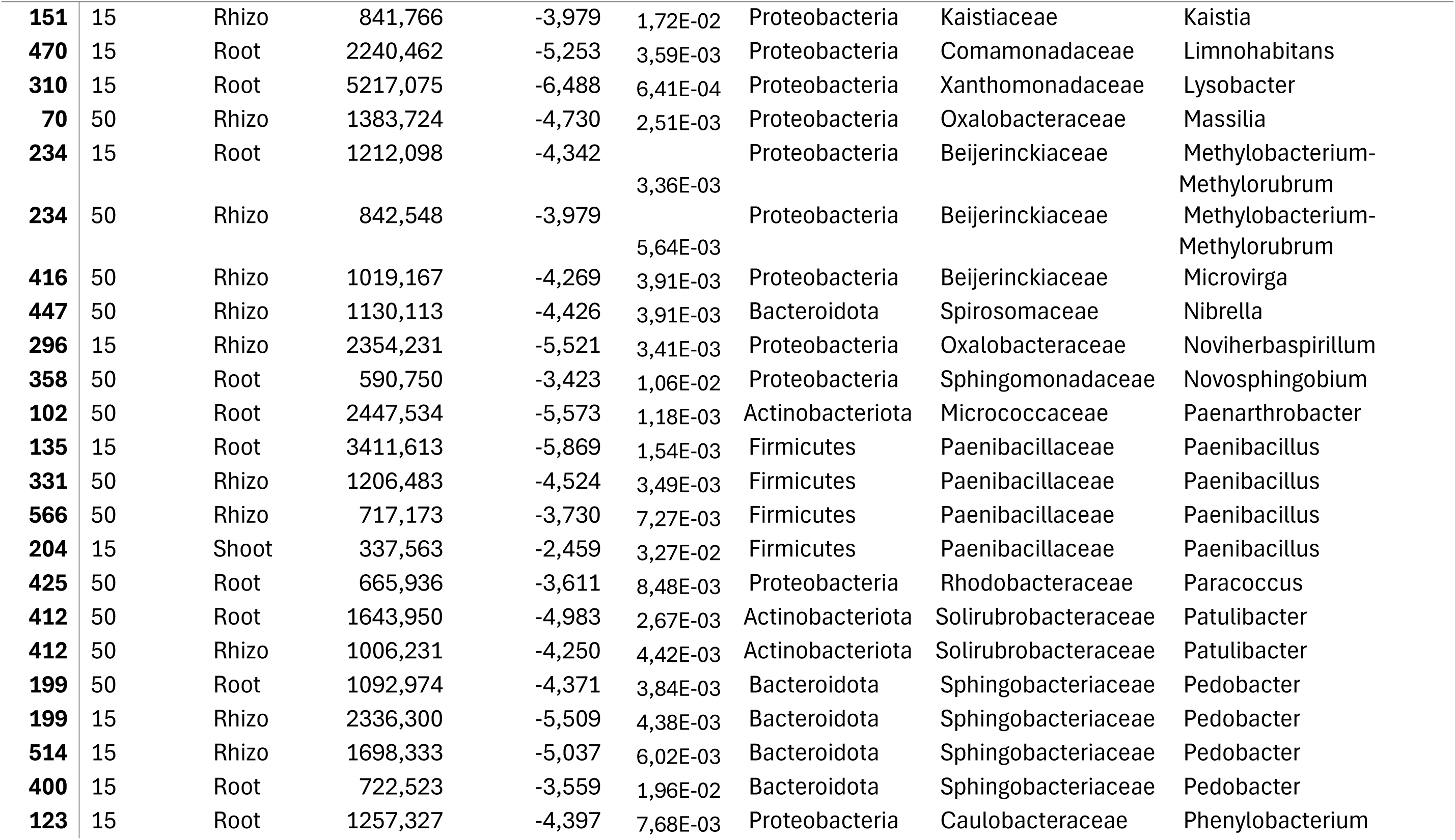
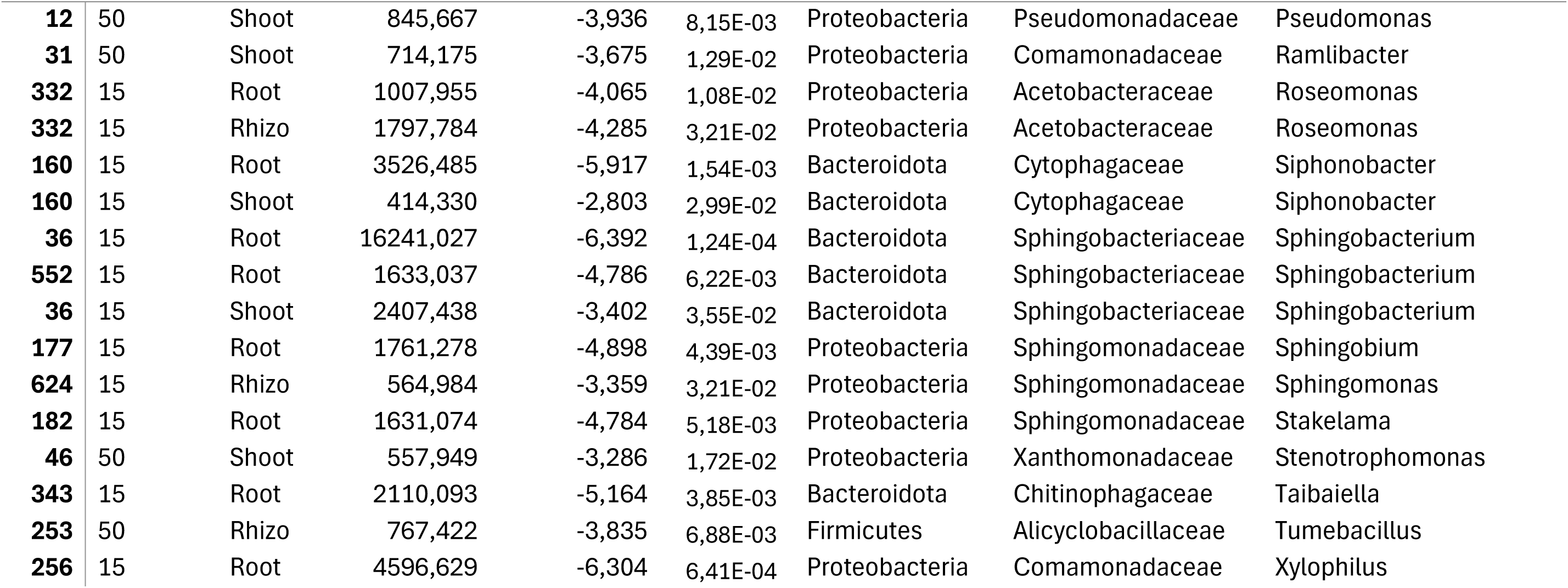
Bacterial ASVs negatively affected by the inoculation.

**Table S3:** Fungal ASVs positively affected by the inoculation.

| ASV | SWC | COMPART | BASEMEAN | LOG2FOLD<br>CHANGE | PADJ | PHYLUM | FAMILY | GENUS |
| --- | --- | --- | --- | --- | --- | --- | --- | --- |
| **71** | 50 | Rhizo | 2012,750 | 6,792 | 1,16E-03 | Ascomycota | SordarialesfamIncertaesedis | Staphylotrichum |
| **64** | 15 | Rhizo | 3475,857 | 7,394 | 8,47E-03 | Basidiomycota | Trichosporonaceae | Cutaneotrichosporon |
| **37** | 15 | Rhizo | 2316,857 | 6,806 | 9,33E-03 | Ascomycota | Aspergillaceae | Aspergillus |
| **71** | 50 | Root | 598,875 | 5,012 | 9,39E-03 | Ascomycota | SordarialesfamIncertaesedis | Staphylotrichum |
| **25** | 15 | Rhizo | 2214,571 | 6,740 | 1,00E-02 | Ascomycota | NA | NA |
| **7** | 50 | Rhizo | 622,750 | 5,070 | 1,03E-02 | Ascomycota | Pleosporaceae | Bipolaris |
| **23** | 15 | Root | 15467,500 | 7,214 | 1,13E-02 | Ascomycota | HypocrealesfamIncertaesedis | Sarocladium |
| **57** | 15 | Shoot | 3244,250 | 5,773 | 1,88E-02 | Ascomycota | Cladosporiaceae | Cladosporium |
| **147** | 50 | Shoot | 401,375 | 4,413 | 2,30E-02 | Basidiomycota | Punctulariaceae | Punctularia |
| **123** | 50 | Rhizo | 338,000 | 4,152 | 2,84E-02 | Ascomycota | Aspergillaceae | Aspergillus |
| **39** | 50 | Rhizo | 233,875 | 3,584 | 4,52E-02 | Ascomycota | Aspergillaceae | Aspergillus |
| **16** | 15 | Rhizo | 53557,286 | 11,346 | 1,89E-10 | Ascomycota | Aspergillaceae | Penicillium |
| **16** | 15 | Root | 2444,000 | 7,074 | 3,23E-06 | Ascomycota | Aspergillaceae | Penicillium |
| **8** | 15 | Rhizo | 53580,857 | 7,778 | 3,28E-05 | Ascomycota | Nectriaceae | Gibberella |
| **16** | 50 | Rhizo | 9642,875 | 9,063 | 3,48E-05 | Ascomycota | Aspergillaceae | Penicillium |
| **11** | 50 | Root | 10712,000 | 9,215 | 3,51E-07 | Ascomycota | Aspergillaceae | Penicillium |
| **16** | 50 | Shoot | 7953,000 | 8,784 | 7,77E-07 | Ascomycota | Aspergillaceae | Penicillium |

**Table S4:** Fungal ASVs negatively affected by the inoculation.

| ASV | %SWHC | COMP. | BASEMEAN | LOG2FOLD<br>CHANGE | PADJ | PHYLUM | FAMILY | GENUS |
| --- | --- | --- | --- | --- | --- | --- | --- | --- |
| **35** | 50 | Shoot | 2217,500 | -6,933 | 1,94E-04 | Ascomycota | Nectriaceae | Gibberella |
| **35** | 50 | Rhizo | 2140,625 | -6,882 | 8,45E-04 | Ascomycota | Nectriaceae | Gibberella |
| **28** | 50 | Root | 1608,125 | -6,465 | 1,06E-03 | Ascomycota | Chaetomiaceae | Humicola |
| **26** | 50 | Shoot | 1783,375 | -6,616 | 1,13E-03 | Ascomycota | Nectriaceae | Gibberella |
| **33** | 50 | Shoot | 1763,750 | -6,600 | 1,13E-03 | Ascomycota | Nectriaceae | Gibberella |
| **95** | 50 | Shoot | 859,250 | -5,546 | 3,73E-03 | Ascomycota | Aspergillaceae | Penicillium |
| **1** | 50 | Root | 110460,375 | -3,842 | 4,88E-02 | Ascomycota | Chaetomiaceae | Ovatospora |
| **35** | 15 | Shoot | 5399,875 | -8,224 | 1,97E-06 | Ascomycota | Nectriaceae | Gibberella |
| **26** | 50 | Root | 32560,125 | -10,820 | 2,03E-07 | Ascomycota | Nectriaceae | Gibberella |
| **2** | 15 | Root | 47153,500 | -9,441 | 3,19E-05 | Ascomycota |  | NA |
| **49** | 15 | Root | 12933,625 | -9,487 | 3,19E-05 | Ascomycota | Nectriaceae | Gibberella |
| **37** | 50 | Root | 5620,250 | -8,282 | 3,90E-05 | Ascomycota | Aspergillaceae | Aspergillus |
| **37** | 50 | Rhizo | 6201,500 | -8,424 | 4,10E-05 | Ascomycota | Aspergillaceae | Aspergillus |

**Table S5.** The effects of irrigation, plant compartment and inoculation on the dominant genera found in the amplicon sequencing datasets based on Kruskal-Wallis test with Benjamini-Hochberg Adjusted p-values.

|  | Compartment (C) | inoculation (I) | SWC | block |
| --- | --- | --- | --- | --- |
| **<u>Bacteria</u>** |  |  |  |  |
| *Flavobacterium* | **P=0.008\*\***<br>**P(adj)=0.028\*** | P=0.749<br>P(adj)=0.873 | **P=0.001\*\***<br>**P(adj)=0.007\*\*** | **P=0.070.**<br>P(adj)=0.148 |
| *Klebsiella* | P=0.849<br>P(adj)=0.849 | **P=0.0003\*\*\***<br>**P(adj)=0.002\*\*** | P=0.966<br>P(adj)=1 | P=0.667<br>P(adj)=0.667 |
| *Bacillus* | **P=0.063.**<br>**P(adj)=0.091.** | P=0.456<br>P(adj)=0.798 | P=0.394<br>P(adj)=1 | **P=0.076.**<br>P(adj)=0.138 |
| *Paenibacillus* | **P =0.0002\*\*\***<br>**P(adj)=0.001\*** | P= 0.209<br>P(adj)=0.646 | P=0.609<br>P(adj)=1 | P=0.537<br>P(adj)=0.626 |
| *Stenotrophomonas* | **P=0.068.**<br>**P(adj)=0.091.** | P= 1<br>P(adj)=1 | P =0.670<br>P(adj)=1 | **P=0.084.**<br>P(adj)=0.138 |
| *Sphingobacterium* | **P =0.039\***<br>**P(adj)=0.091.** | P=0.686<br>P(adj)=0.873 | P =1<br>P(adj)=1 | **P=0.009\*\***<br>**P(adj)=0.063.** |
| **<u>Fungi</u>** |  |  |  |  |
| *Zopfiella* | P=0.570<br>P(adj)=0.57 | **P=0.004\*\***<br>**P(adj)=0.036\*** | P=0.147<br>P(adj)=0.436 | **P=0.070**<br>P(adj)=0.171 |
| *Epicoccum* | **P=0.003\*\***<br>**P(adj)=0.009\*\*** | P=0.798<br>P(adj)=0.897 | P=0.848<br>P(adj)=0.983 | P=0.45<br>P(adj)=0.578 |
| *Aspergillus* | P=0.318<br>P(adj)=0.357 | P=0.317<br>P(adj)= 0.709 | **P=0.028\***<br>P(adj)=0.252 | **P=0.076.**<br>P(adj)=0.171 |
| *Penicilium* | **P=0.0002\*\*\***<br>**P(adj)=0.001\*\*** | P=0.317<br>P(adj)= 0.709 | P=0.983<br>P(adj)=0.983 | **P=0.097.**<br>P(adj)=0.174 |
| *Humicola* | **P=0.0003\*\*\***<br>**P(adj)=0.001\*\*** | P=0.966<br>P(adj)= 0.966 | P=0.443<br>P(adj)=0.797 | **P=0.066.**<br>P(adj)=0.171 |
| *Papulaspora* | P=0.172<br>P(adj)=0.258 | **P=0.041\***<br>P(adj)= 0.184 | P=0.551<br>P(adj)=0.826 | P=0.707<br>P(adj)=0.707 |

\*\*\*: P < 0.001, \*\*: 0.001 < P < 0.01, \*:0.01 < P < 0.05, .: 0.05 < P < 0.10

## References

1. Farrar K, Bryant D, Cope-Selby N: Understanding and engineering beneficial plant–microbe interactions: plant growth promotion in energy crops. Plant Biotechnology Journal 2014, 12:1193–1206.

2. Quiza L, St-Arnaud M, Yergeau E: Harnessing phytomicrobiome signaling for rhizosphere microbiome engineering. Front Plant Sci 2015, 6.

3. Agoussar A, Yergeau E: Engineering the plant microbiota in the context of the theory of ecological communities. Current Opinion in Biotechnology 2021, 70:220–225.

4. Emami S, Alikhani HA, Pourbabaee AA, Etesami H, Motasharezadeh B, Sarmadian F: Consortium of endophyte and rhizosphere phosphate solubilizing bacteria improves phosphorous use efficiency in wheat cultivars in phosphorus deficient soils. Rhizosphere 2020, 14:100196.

5. Giri S, Shitut S, Kost C: Harnessing ecological and evolutionary principles to guide the design of microbial production consortia. Current Opinion in Biotechnology 2020, 62:228–238.

6. Hone H, Mann R, Yang G, Kaur J, Tannenbaum I, Li T, Spangenberg G, Sawbridge T: Profiling, isolation and characterisation of beneficial microbes from the seed microbiomes of drought tolerant wheat. Sci Rep 2021, 11:11916.

7. Azarbad H, Constant P, Giard-Laliberté C, Bainard LD, Yergeau E: Water stress history and wheat genotype modulate rhizosphere microbial response to drought. Soil Biology and Biochemistry 2018, 126:228–236.

8. Azarbad H, Tremblay J, Giard-Laliberté C, Bainard LD, Yergeau E: Four decades of soil water stress history together with host genotype constrain the response of the wheat microbiome to soil moisture. FEMS Microbiology Ecology 2020, 96.

9. Giard-Laliberté C, Azarbad H, Tremblay J, Bainard L, Yergeau É: A water stress-adapted inoculum affects rhizosphere fungi, but not bacteria nor wheat. FEMS Microbiol Ecol 2019, doi:10.1093/femsec/fiz080.

10. Agoussar A, Azarbad H, Tremblay J, Yergeau É: The resistance of the wheat microbial community to water stress is more influenced by plant compartment than reduced water availability. FEMS Microbiol Ecol 2021, 97.

11. Azarbad H, Bainard LD, Agoussar A, Tremblay J, Yergeau E: The response of wheat and its microbiome to contemporary and historical water stress in a field experiment. ISME COMMUN 2022, 2:1–9.

12. Pande PM, Azarbad H, Tremblay J, St-Arnaud M, Yergeau E: Metatranscriptomic response of the wheat holobiont to decreasing soil water content. ISME COMMUN 2023, 3:1–13.

13. Schmidt RL, Azarbad H, Bainard L, Tremblay J, Yergeau E: Intermittent water stress favors microbial traits that better help wheat under drought. ISME Communications 2024, 4:ycae074.

14. Preece C, Peñuelas J: Rhizodeposition under drought and consequences for soil communities and ecosystem resilience. Plant and Soil 2016, 409:1–17.

15. Mallon CA, Le Roux X, van Doorn GS, Dini-Andreote F, Poly F, Salles JF: The impact of failure: unsuccessful bacterial invasions steer the soil microbial community away from the invader’s niche. ISME J 2018, 12:728–741.

16. Mawarda PC, Le Roux X, Dirk van Elsas J, Salles JF: Deliberate introduction of invisible invaders: A critical appraisal of the impact of microbial inoculants on soil microbial communities. Soil Biology and Biochemistry 2020, 148:107874.

17. Taghavi S, Barac T, Greenberg B, Borremans B, Vangronsveld J, Lelie D van der: Horizontal Gene Transfer to Endogenous Endophytic Bacteria from Poplar Improves Phytoremediation of Toluene. Appl Environ Microbiol 2005, 71:8500– 8505.

18. Barcellos FG, Menna P, da Silva Batista JS, Hungria M: Evidence of Horizontal Transfer of Symbiotic Genes from a Bradyrhizobium japonicum Inoculant Strain to Indigenous Diazotrophs Sinorhizobium (Ensifer) fredii and Bradyrhizobium elkanii in a Brazilian Savannah Soil. Applied and Environmental Microbiology 2007, 73:2635–2643.

19. Ling J, Wang H, Wu P, Li T, Tang Y, Naseer N, Zheng H, Masson-Boivin C, Zhong Z, Zhu J: Plant nodulation inducers enhance horizontal gene transfer of Azorhizobium caulinodans symbiosis island. Proceedings of the National Academy of Sciences 2016, 113:13875–13880.

20. Figueiredo MVB, Burity HA, Martínez CR, Chanway CP: Alleviation of drought stress in the common bean (Phaseolus vulgaris L.) by co-inoculation with Paenibacillus polymyxa and Rhizobium tropici. Applied Soil Ecology 2008, 40:182–188.

21. Riaz U, Murtaza G, Anum W, Samreen T, Sarfraz M, Nazir MZ: Plant Growth-Promoting Rhizobacteria (PGPR) as Biofertilizers and Biopesticides. In Microbiota and Biofertilizers: A Sustainable Continuum for Plant and Soil Health. Edited by Hakeem KR, Dar GH, Mehmood MA, Bhat RA. Springer International Publishing; 2021:181–196.

22. Yadav VK, Bhagat N, Sharma SK: Modulation in Plant Growth and Drought Tolerance of Wheat Crop upon Inoculation of Drought-tolerant-Bacillus Species Isolated from Hot Arid Soil of India. Journal of Pure and Applied Microbiology 2022, 16:246–263.

23. Babalola OO: Beneficial bacteria of agricultural importance. Biotechnol Lett 2010, 32:1559–1570.

24. Camaille M, Fabre N, Clément C, Ait Barka E: Advances in Wheat Physiology in Response to Drought and the Role of Plant Growth Promoting Rhizobacteria to Trigger Drought Tolerance. Microorganisms 2021, 9:687.

25. Sandhya V, Z AS, Grover M, Reddy G, Venkateswarlu B: Alleviation of drought stress effects in sunflower seedlings by the exopolysaccharides producing Pseudomonas putida strain GAP-P45. Biol Fertil Soils 2009, 46:17–26.

26. Redman RS, Kim YO, Woodward CJDA, Greer C, Espino L, Doty SL, Rodriguez RJ: Increased Fitness of Rice Plants to Abiotic Stress Via Habitat Adapted Symbiosis: A Strategy for Mitigating Impacts of Climate Change. PLOS ONE 2011, 6:e14823.

27. Timmusk S, Paalme V, Pavlicek T, Bergquist J, Vangala A, Danilas T, Nevo E: Bacterial Distribution in the Rhizosphere of Wild Barley under Contrasting Microclimates. PLOS ONE 2011, 6:e17968.

28. Glick BR: Bacteria with ACC deaminase can promote plant growth and help to feed the world. Microbiological Research 2014, 169:30–39.

29. Defez R, Andreozzi A, Dickinson M, Charlton A, Tadini L, Pesaresi P, Bianco C: Improved Drought Stress Response in Alfalfa Plants Nodulated by an IAA Over-producing Rhizobium Strain. Front Microbiol 2017, 8.

30. Madkour MA, Smith LT, Smith GM: Preferential Osmolyte Accumulation: a Mechanism of Osmotic Stress Adaptation in Diazotrophic Bacteria. Applied and Environmental Microbiology 1990, 56:2876–2881.

31. Gagné-Bourque F, Bertrand A, Claessens A, Aliferis KA, Jabaji S: Alleviation of Drought Stress and Metabolic Changes in Timothy (Phleum pratense L.) Colonized with Bacillus subtilis B26. Frontiers in Plant Science 2016, 7.

32. Vílchez JI, Niehaus K, Dowling DN, González-López J, Manzanera M: Protection of Pepper Plants from Drought by Microbacterium sp. 3J1 by Modulation of the Plant’s Glutamine and α-ketoglutarate Content: A Comparative Metabolomics Approach. Front Microbiol 2018, 9.

33. Khan N, Bano A, Rahman MA, Guo J, Kang Z, Babar MA: Comparative Physiological and Metabolic Analysis Reveals a Complex Mechanism Involved in Drought Tolerance in Chickpea (Cicer arietinum L.) Induced by PGPR and PGRs. Sci Rep 2019, 9:2097.

34. Paul D, Nair S: Stress adaptations in a Plant Growth Promoting Rhizobacterium (PGPR) with increasing salinity in the coastal agricultural soils. Journal of Basic Microbiology 2008, 48:378–384.

35. Hubbard M: Fungal endophytes that confer heat and drought tolerance to wheat. 2012,

36. Hubbard M, Germida JJ, Vujanovic V: Fungal endophyte colonization coincides with altered DNA methylation in drought-stressed wheat seedlings. Can J Plant Sci 2014, 94:223–234.

37. Zolla G, Badri DV, Bakker MG, Manter DK, Vivanco JM: Soil microbiomes vary in their ability to confer drought tolerance to Arabidopsis. Applied Soil Ecology 2013, 68:1–9.

38. Agoussar A, Azarbad H, Tremblay J, Yergeau É: The resistance of the wheat microbial community to water stress is more influenced by plant compartment than reduced water availability. FEMS Microbiology Ecology 2021, 97:fiab149.

39. Eng A, Borenstein E: Microbial community design: methods, applications, and opportunities. Current Opinion in Biotechnology 2019, 58:117–128.

40. Khan IA, Ahmad S, Ayub N: Response of Oat (Avena sativa) to Inoculation with Vesicular Arbuscular Mycorrhizae (VAM) in the Presence of Phosphorus. Asian Journal of Plant Sciences 2003,

41. Maymon M, Martínez-Hidalgo P, Tran S, Ice T, Craemer K, Anbarchian T, Sung T, Hwang L, Chou M, Fujishige N, et al.: Mining the phytomicrobiome to understand how bacterial coinoculations enhance plant growth. Frontiers in Plant Science 2015, 6.

42. Riis V, Lorbeer H, Babel W: Extraction of microorganisms from soil: evaluation of the efficiency by counting methods and activity measurements. Soil Biology and Biochemistry 1998, 30:1573–1581.

43. Tardif S, Yergeau É, Tremblay J, Legendre P, Whyte LG, Greer CW: The Willow Microbiome Is Influenced by Soil Petroleum-Hydrocarbon Concentration with Plant Compartment-Specific Effects. Front Microbiol 2016, 7.

44. Maynard DN, Hochmuth George J: Knott’s Handbook for Vegetable Growers. John Wiley & Sons, Inc.; 2007.

45. Tremblay J, Yergeau E: Systematic processing of ribosomal RNA gene amplicon sequencing data. GigaScience 2019, 8.

46. Edgar RC, Haas BJ, Clemente JC, Quince C, Knight R: UCHIME improves sensitivity and speed of chimera detection. Bioinformatics 2011, 27:2194–2200.

47. Quast C, Pruesse E, Yilmaz P, Gerken J, Schweer T, Yarza P, Peplies J, Glöckner FO: The SILVA ribosomal RNA gene database project: improved data processing and web-based tools. Nucleic Acids Research 2013, 41:D590–D596.

48. David AS, Thapa-Magar KB, Menges ES, Searcy CA, Afkhami ME: Do plant– microbe interactions support the Stress Gradient Hypothesis? Ecology 2020, 101:e03081.

49. Bell TH, Stefani FOP, Abram K, Champagne J, Yergeau E, Hijri M, St-Arnaud M: A Diverse Soil Microbiome Degrades More Crude Oil than Specialized Bacterial Assemblages Obtained in Culture. Appl Environ Microbiol 2016, 82:5530–5541.

50. Yergeau E, Bell TH, Champagne J, Maynard C, Tardif S, Tremblay J, Greer CW: Transplanting Soil Microbiomes Leads to Lasting Effects on Willow Growth, but not on the Rhizosphere Microbiome. Front Microbiol 2015, 6.

51. Ravel C, Courty C, Coudret A, Charmet G: Beneficial effects of Neotyphodium lolii on the growth and the water status in perennial ryegrass cultivated under nitrogen deficiency or drought stress. Agronomie 1997, 17:173–181.

52. Kavroulakis N, Doupis G, Papadakis IE, Ehaliotis C, Papadopoulou KK: Tolerance of tomato plants to water stress is improved by the root endophyte Fusarium solani FsK. Rhizosphere 2018, 6:77–85.

53. Wang Q, Garrity GM, Tiedje JM, Cole JR: Naïve Bayesian Classifier for Rapid Assignment of rRNA Sequences into the New Bacterial Taxonomy. Appl Environ Microbiol 2007, 73:5261–5267.

54. Schimel J, Balser TC, Wallenstein M: MICROBIAL STRESS-RESPONSE PHYSIOLOGY AND ITS IMPLICATIONS FOR ECOSYSTEM FUNCTION. Ecology 2007, 88:1386–1394.

55. Vries FT de, Griffiths RI, Knight CG, Nicolitch O, Williams A: Harnessing rhizosphere microbiomes for drought-resilient crop production. Science 2020, 368:270–274.

56. Scholz A, Stahl J, de Berardinis V, Müller V, Averhoff B: Osmotic stress response in Acinetobacter baylyi: identification of a glycine–betaine biosynthesis pathway and regulation of osmoadaptive choline uptake and glycine–betaine synthesis through a choline-responsive BetI repressor. Environmental Microbiology Reports 2016, 8:316–322.

57. Guo J, Deng X, Zhang Y, Song S, Zhao T, Zhu D, Cao S, Baryshnikov PI, Cao G, Blair HT, et al.: The Flagellar Transcriptional Regulator FtcR Controls Brucella melitensis 16M Biofilm Formation via a betI-Mediated Pathway in Response to Hyperosmotic Stress. International Journal of Molecular Sciences 2022, 23:9905.

58. Subhadra B, Surendran S, Lim BR, Yim JS, Kim DH, Woo K, Kim H-J, Oh MH, Choi CH: The osmotic stress response operon betIBA is under the functional regulation of BetI and the quorum-sensing regulator AnoR in Acinetobacter nosocomialis. J Microbiol 2020, 58:519–529.

59. Scott JM, Haldenwang WG: Obg, an Essential GTP Binding Protein ofBacillus subtilis, Is Necessary for Stress Activation of Transcription Factor ςB. Journal of Bacteriology 1999, 181:4653–4660.

60. Kim JF, Jeong H, Park S-Y, Kim S-B, Park YK, Choi S-K, Ryu C-M, Hur C-G, Ghim S-Y, Oh TK, et al.: Genome Sequence of the Polymyxin-Producing Plant-Probiotic Rhizobacterium Paenibacillus polymyxa E681. Journal of Bacteriology 2010, 192:6103–6104.

61. Mahapatra S, Yadav R, Ramakrishna W: Bacillus subtilis impact on plant growth, soil health and environment: Dr. Jekyll and Mr. Hyde. Journal of Applied Microbiology 2022, 132:3543–3562.

62. Jeong H, Choi S-K, Ryu C-M, Park S-H: Chronicle of a Soil Bacterium: Paenibacillus polymyxa E681 as a Tiny Guardian of Plant and Human Health. Frontiers in Microbiology 2019, 10.

63. Boonmak C, Kettongruang S, Buranathong B, Morikawa M, Duangmal K: Duckweed-associated bacteria as plant growth-promotor to enhance growth of Spirodela polyrhiza in wastewater effluent from a poultry farm. Arch Microbiol 2023, 206:43.

64. Giongo A, Arnhold J, Grunwald D, Smalla K, Braun-Kiewnick A: Soil depths and microhabitats shape soil and root-associated bacterial and archaeal communities more than crop rotation in wheat. Front Microbiomes 2024, 3.

65. Mataranyika PN, Bez C, Venturi V, Chimwamurombe PM, Uzabakiriho JD: Rhizospheric, seed, and root endophytic-associated bacteria of drought-tolerant legumes grown in arid soils of Namibia. Heliyon 2024, 10.

66. Gilbert J, Fernando WGD: Epidemiology and biological control of Gibberella zeae / Fusarium graminearum. Canadian Journal of Plant Pathology 2004, 26:464– 472.

67. Cumagun CJR, Bowden RL, Jurgenson JE, Leslie JF, Miedaner T: Genetic Mapping of Pathogenicity and Aggressiveness of Gibberella zeae (Fusarium graminearum) Toward Wheat. Phytopathology® 2004, 94:520–526.

68. Abd El Daim IA, Häggblom P, Karlsson M, Stenström E, Timmusk S: Paenibacillus polymyxa A26 Sfp-type PPTase inactivation limits bacterial antagonism against Fusarium graminearum but not of F. culmorum in kernel assay. Frontiers in Plant Science 2015, 6.

69. Saxena AK, Kumar M, Chakdar H, Anuroopa N, Bagyaraj DJ: Bacillus species in soil as a natural resource for plant health and nutrition. J Appl Microbiol 2020, 128:1583–1594.

70. Rashid U, Yasmin H, Hassan MN, Naz R, Nosheen A, Sajjad M, Ilyas N, Keyani R, Jabeen Z, Mumtaz S, et al.: Drought-tolerant Bacillus megaterium isolated from semi-arid conditions induces systemic tolerance of wheat under drought conditions. Plant Cell Rep 2022, 41:549–569.

71. Xue H, Tu Y, Ma T, Jiang N, Piao C, Li Y: Taxonomic Study of Three Novel Paenibacillus Species with Cold-Adapted Plant Growth-Promoting Capacities Isolated from Root of Larix gmelinii. Microorganisms 2023, 11:130.

72. Singh RP, Jha PN: The PGPR Stenotrophomonas maltophilia SBP-9 Augments Resistance against Biotic and Abiotic Stress in Wheat Plants. Frontiers in Microbiology 2017, 8.

73. Kasim WA, Osman MEH, Omar MN, Salama S: Enhancement of drought tolerance in Triticum aestivum L. seedlings using Azospirillum brasilense NO40 and Stenotrophomonas maltophilia B11. Bulletin of the National Research Centre 2021, 45:95.

74. Hagaggi NShA, Abdul-Raouf UM: Drought-tolerant Sphingobacterium changzhouense Alv associated with Aloe vera mediates drought tolerance in maize (Zea mays). World J Microbiol Biotechnol 2022, 38:248.

75. Kour D, Rana KL, Yadav AN, Sheikh I, Kumar V, Dhaliwal HS, Saxena AK: Amelioration of drought stress in Foxtail millet (Setaria italica L.) by P-solubilizing drought-tolerant microbes with multifarious plant growth promoting attributes. Environmental Sustainability 2020, 3:23–34.

76. Miranda V, Silva-Castro GA, Ruiz-Lozano JM, Fracchia S, García-Romera I: Fungal Endophytes Enhance Wheat and Tomato Drought Tolerance in Terms of Plant Growth and Biochemical Parameters. Journal of Fungi 2023, 9:384.

77. Kour D, Rana KL, Kaur T, Sheikh I, Yadav AN, Kumar V, Dhaliwal HS, Saxena AK: Microbe-mediated alleviation of drought stress and acquisition of phosphorus in great millet (Sorghum bicolour L.) by drought-adaptive and phosphorus-solubilizing microbes. Biocatalysis and Agricultural Biotechnology 2020, 23:101501.

